# Systematic identification and characterization of regulators of aryl hydrocarbon receptor signaling

**DOI:** 10.1101/2025.09.15.676391

**Authors:** Manasvi Verma, Kushaal Desai, Yufang Ding, Xingren Wang, Minwoo Bae, Seth Rakoff-Nahoum, Emily P. Balskus, Michael A. Fischbach, Marco Jost

## Abstract

The human aryl hydrocarbon receptor (AHR) integrates chemical signals derived from the environment, gut microbes, and endogenous sources to regulate processes ranging from intestinal barrier integrity to xenobiotic detoxification. Despite strong evidence that dysregulation of AHR signaling is a causal factor in metabolic and autoimmune disorders, we currently lack a comprehensive understanding of the factors that regulate AHR activity in human cells. Here, we use genome-scale CRISPR screening to systematically identify regulators of AHR signaling in hepatocytes. The resulting datasets recapitulate the core AHR signaling pathway and identify a large network of regulators. Many of these factors have roles beyond AHR signaling, reflecting that AHR signaling is deeply integrated into human cell biology. We further dissect this network to reveal novel modes of regulation of AHR expression, protein levels, and signaling. For example, we find that the E3 ubiquitin ligase UBR5 sustains AHR signaling by counteracting degradation of ligand-bound AHR. Finally, we identify components of the AHR regulatory network that are specific to cell types and ligands as potential nodes to manipulate AHR signaling in a targeted manner for therapeutic benefit. Overall, our results define the regulatory network that underpins AHR activation, with implications for our understanding of host-microbe interactions and integrative chemosensation and the etiology of metabolic and inflammatory disorders.

## Background

The human aryl hydrocarbon receptor (AHR) is a ligand-activated transcription factor that mediates responses to a large panel of chemicals derived from the environment, the diet, gut microbes, and endogenous sources (1, 2). AHR was initially discovered in the 1970s as the receptor that recognizes and mediates the biological effects of synthetic environmental toxins, such as halogenated and polycyclic aromatic hydrocarbons (3). Subsequent discoveries of developmental defects in *Ahr* knockout mice (4, 5) and findings that AHR is broadly expressed across the body spurred broader investigation of the physiological roles of AHR, which revealed roles of AHR signaling in many additional processes. Among others, AHR activation in hepatocytes results in upregulation of enzymes involved in xenobiotic and drug metabolism, such as cytochrome P450s (6-8); AHR activation in naïve T-cells promotes differentiation of Th17 cells (9,10), which can exacerbate autoimmunity in mouse models (9); AHR activation in intestinal lymphocytes promotes intestinal barrier integrity (11,12); and AHR activation in some cancer cells promotes proliferation and suppresses anti-tumor immunity (13-15). Complementary genetic studies in the human population have linked mutations in regulatory regions of AHR to multiple types of skin cancer (16) and loss-of-function mutations in AHR to retinitis pigmentosa and foveal hypoplasia (17-19) Overall, both overactivation and loss of AHR signaling have been implicated in inflammatory and metabolic diseases as well as cancer (9-26).

Parallel efforts to understand how AHR activity is regulated have established the core pathway of AHR signaling (Figure 1A) (1,27). In the absence of a ligand, AHR is held in an inactive form in the cytosol in a complex with HSP90, p23, and AHR interacting protein (AIP) (28). Ligand binding triggers nuclear translocation of AHR and heterodimerization with AHR nuclear translocator (ARNT, also known as HIF-1β) (29, 30). The AHR-ARNT complex binds to DNA at specific motifs known as AHR response elements (AREs, also called xenobiotic response elements) and activates transcription of target genes (29-31), the identities of which vary by cell type. In most cell types, these target genes include at least one of the three known negative regulators of AHR signaling: (i) *CYP1A1*, which encodes a cytochrome P450 that degrades some known AHR ligands (32); (ii) *TIPARP* (TCDD-inducible poly-ADP-ribose-polymerase), which encodes an ADP-ribosyltransferase that targets AHR for degradation by catalyzing mono-ADP-ribosylation of AHR (33, 34); and (iii) *AHRR* (AHR repressor), which encodes a repressor that competes with ARNT for binding to AHR (35). Together, these three factors mediate negative feedback regulation of AHR signaling (10, 12, 33-35).

**Figure 1.**
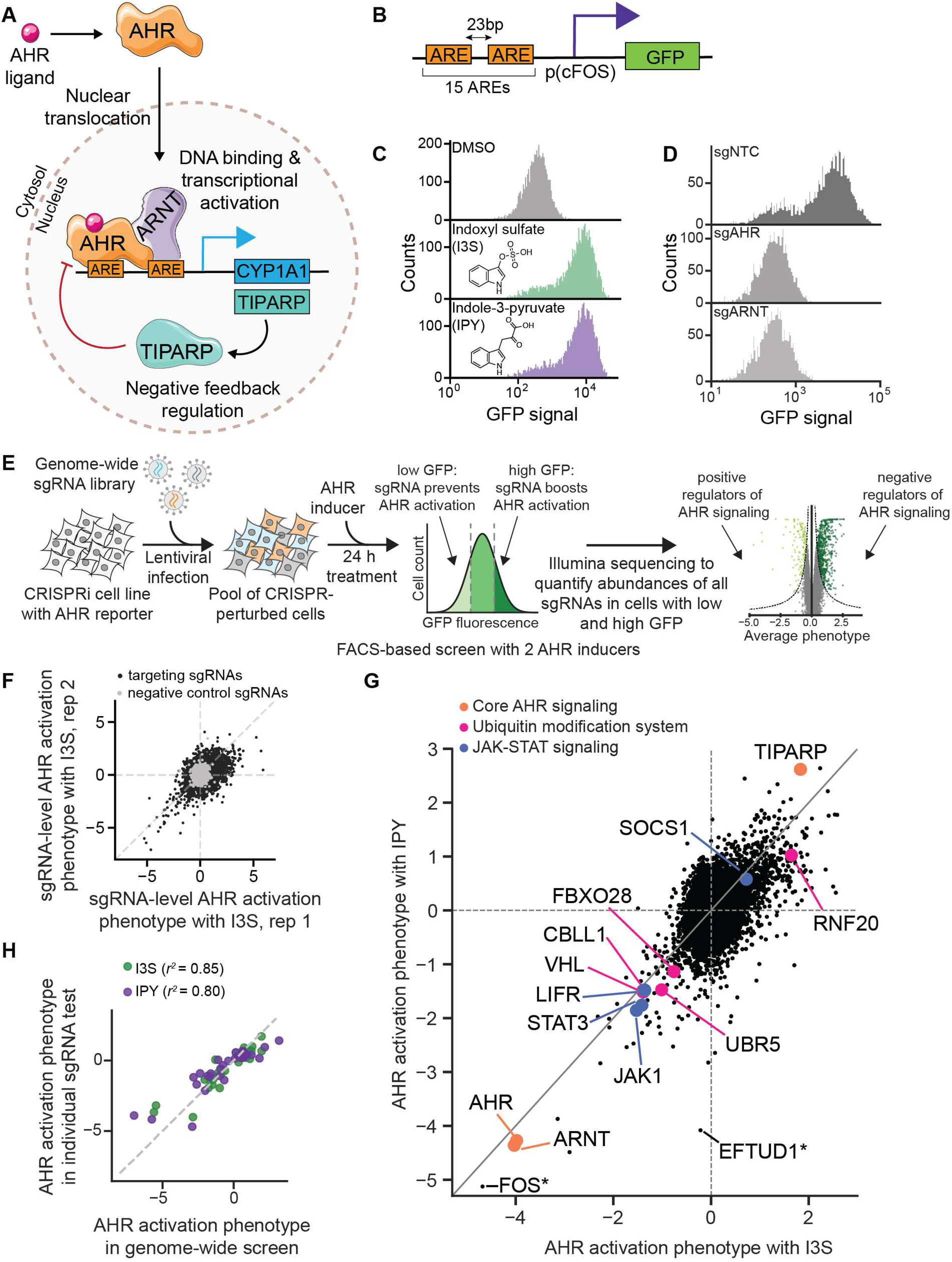
Genome-scale CRISPRi screens reveal modulators of AHR signaling. (A) Overview of the core AHR signaling pathway. (B) Schematic of GFP-based AHR reporter. (C) GFP fluorescence distribution of monoclonal HepG2 reporter cells (cMV011) treated with 100 µM indoxyl sulfate (I3S) or 100 µM indole-3-pyruvate (IPY) for 24 h, as measured using flow cytometry. (D) GFP fluorescence of HepG2 reporter cells transduced with the indicated sgRNAs and treated with 100 µM IPY for 24 h. Data are representative of two independent transductions. sgNTC is a non-targeting control sgRNA. (E) Schematic of genome-scale CRISPRi screen. (F) AHR activation phenotypes of all sgRNAs from two biological replicates of the CRISPRi screen treated with I3S. Non-targeting control sgRNAs are shown in gray and targeting sgRNAs in black. (G) AHR activation phenotypes for all genes. Categories of genes highlighted: core AHR signaling genes (orange), genes involved in ubiquitin-mediated protein degradation (pink), and genes that are components of the JAK-STAT signaling pathway (blue). FOS (marked with *) is the strongest hit, but the phenotypes likely result from sgRNAs targeting the p(cFos) promoter of the AHR reporter, as further shown in Figure S1C. EFTUD1 (marked with *) shows the strongest difference between the two inducers but is essential. (H) Correlation between AHR activation phenotypes in the CRISPRi screen and a subsequent validation experiment performed by individually transducing sgRNAs and measuring their GFP reporter phenotypes using flow cytometry. *r*^2^ is squared Pearson correlation coefficient. Phenotypes are average of two independently transduced replicates except for the indole-3-pyruvate screen phenotypes.

Beyond this core pathway, however, it has become clear that AHR signaling is regulated by many additional factors, the identities of which can vary across cell types and ligands (1, 13, 36-42). For example, in murine liver cells, cinnabarinic acid (CA), but not 2,3,7,8-Tetrachlorodibenzo-p-dioxin (TCDD), recruits the epigenetic regulators Mta2, Atf2, and Dot1l to initiate transcription of Stanniocalcin-2, a CA-specific AHR target gene (40, 41). Further, crosstalk between AHR and multiple additional transcription factors, including NF-κB, RB1, E2F1, and ESR1, has been described in some cell types but has not been evaluated across multiple cell types and ligands (13, 43). We currently do not have a comprehensive understanding of the regulation of AHR signaling for any single cell type or ligand, let alone across the diverse cell types in which AHR is active and the broad range of ligands to which AHR responds.

The development of CRISPR-mediated genetics has now made it possible to identify regulators of signaling pathways in human cells in a systematic manner (42-46). Here, we develop an optimized reporter system for AHR signaling, with which we conduct genome-scale CRISPR interference (CRISPRi) screens for regulators of AHR activity in human cells. The resulting datasets reveal a large network of regulators, the composition of which varies across ligands and cell types. Through mechanistic interrogation, we place individual regulators at different steps of the signaling pathway from regulation of AHR expression to regulation of AHR signaling. Our results provide a comprehensive picture of the regulatory network that underpins AHR signaling and provide a foundation to target AHR signaling in a specific manner for therapeutic benefit.

## Results

### Genome-wide CRISPRi screens reveal modulators of AHR signaling

To enable pooled genetic screens for regulators of AHR signaling, we first sought to establish a fluorescent reporter for AHR activity that has high sensitivity and dynamic range and can be stably integrated into human cells. Although some fluorescent reporters for AHR had been reported (23, 47), they typically had limited dynamic range. We designed a lentiviral reporter in which expression of EGFP is driven by a minimal promoter with a series of AHR response elements, akin to similar designs for other transcription factors (42-46). This construct reports on AHR activity at endogenous expression levels using cognate DNA binding motifs of AHR. We iteratively optimized several design elements including the number of AHR response elements (AREs), the spacing between them, and the identity of the minimal promoter (Figure 1B, S1A). Our final reporter contains 15 AREs spaced 23 bp apart followed by a minimal cFos promoter (p(cFos)). We integrated this reporter into an existing HepG2 (human hepatocellular carcinoma) cell line that stably expresses the Zim3-dCas9 effector for CRISPR-mediated knockdown (CRISPR interference, CRISPRi) (48). A resulting monoclonal reporter cell line demonstrates a strong and homogenous increase in GFP signal after treatment with 100 µM indoxyl sulfate (I3S) or 100 µM indole-3-pyruvate (IPY), two well-characterized AHR inducers, for 24 h, as measured by flow cytometry (Figure 1C). CRISPRi-mediated knockdown of AHR or ARNT abolished reporter induction by IPY (Figure 1D), confirming that the cell line is specifically responsive to AHR.

To identify regulators of AHR activity, we next conducted genome-scale, FACS-based CRISPRi screens for genes whose knockdown decreases or increases AHR activity (Figure 1E). We conducted these screens with two AHR inducers to enable comparison of regulators across inducers: I3S, a well-characterized AHR agonist that is derived from the gut microbiome (49), and IPY, a precursor to a strong agonist (50-52) that is produced by both endogenous and microbial metabolism. Briefly, we transduced the CRISPRi AHR reporter cell line in duplicate with the hCRISPRi-v2-top5 sgRNA library, which targets 18,905 genes with 5 sgRNAs for each transcription start site (TSS) (53).

We cultured the cells at a coverage of >1000 cells/sgRNA for 10 days, treated them with 100 µM I3S (both replicates) or 100 µM IPY (one replicate) for 24 h, and used FACS to isolate cells with high (top 30%) or low (bottom 30%) GFP signal. Finally, we measured the relative abundance of each sgRNA in each population using Illumina sequencing.

We calculated effect sizes for each sgRNA, which we term AHR activation phenotypes, as the log_2_ ratio of sgRNA counts within the high GFP cells over the low GFP cells, normalized to the median log_2_ ratio of negative control sgRNAs. AHR activation phenotypes of targeting sgRNAs were well-correlated between replicates of the I3S screen whereas those for non-targeting sgRNAs were clustered around 0 (Figure 1F, Table S1). Finally, we quantified gene-level phenotypes by averaging phenotypes of the 3 sgRNAs with the strongest phenotype by absolute value (Table S2).

Gene phenotypes for AHR activation by I3S and IPY correlated well (Pearson *r*^2^ = 0.75 for genes with absolute phenotypes > 1.5), implying that knockdown of most genes affects AHR activation by I3S and IPY similarly (Figure 1G, Table S2). Among the strongest hits are known modulators of AHR activation, including (i) *AHR* itself, (ii) *ARNT*, whose knockdown inhibited AHR activation, and (iii) *TIPARP*, whose knockdown enhanced AHR activation (Figure 1A, 1G, S1B). Three sgRNAs targeting *FOS* also led to strong decreases in reporter signal (Figure 1G), but all three sgRNAs have perfect matches within the p(cFos) promoter in the reporter. In contrast, several additional *FOS*-targeting sgRNAs that resulted in strong knockdown of *FOS* did not affect AHR activation (Figure S1C). We thus attribute the effects of these sgRNAs to repression of the reporter rather than knockdown of *FOS*. The remaining hits fall into several broad categories: (i) genes involved in protein homeostasis, including the E3 ligases *UBR5*, *VHL*, *CBLL1, FBXO28*, and *RNF20* (ii) genes in the JAK/STAT signaling pathway, including *JAK1*, *STAT3*, *LIFR*, and *SOCS1*; (iii) multiple other transcription factors, including *HNF1A, FOXA2, NR5A1,* and *NR2F2*; and (iv) general transcriptional machinery, including basal transcription factors (*GTF2A1, GTF2A2, GTF2E1, GTF2F2, GTF2H4,*), histone modifiers (*KMT2A, SETDB1, EHMT2*), and chromatin remodelers (*SMARCA5, BPTF*) (Figure 1G, S1D). Many of these genes had not been implicated in regulation of AHR activation.

Knockdown of a small number of genes affected AHR activation by only one of the inducers, such as *EFTUD1* (now *EFL1*), which encodes a GTPase involved in ribosome assembly (Figure 1G). Further inspection revealed that most of these genes are essential and that AHR activation phenotypes were driven by sgRNAs with low overall counts, which are more subject to experimental noise. We conclude that the inducer-specific phenotypes of these genes are unlikely to be meaningful.

To validate the results of the screen, we measured the effects of 47 targeting sgRNAs and 1 non-targeting control sgRNA in an independent experiment using an internally controlled assay. Briefly, we cloned each sgRNA into a lentiviral vector that also expresses BFP as a marker, transduced each vector in duplicate into our AHR reporter cell line in an arrayed format, and measured GFP induction after treatment with 100 µM I3S or 100 µM IPY for 24 h by flow cytometry. Non-transduced cells in the same well (BFP-negative) served as internal controls for AHR activation. We excluded 15 sgRNAs for which the number of sgRNA-expressing cells at the end point was too low (< 100 cells) to accurately quantify GFP levels. For these sgRNAs, counts of sgRNA-expressing cells decreased rapidly over time, suggesting that the targeted genes are essential (Figure S1E). Reporter phenotypes of the remaining genes matched those of the screen (Figure 1H, S1F). Thus, our genetic screening strategy accurately and reproducibly identified regulators of AHR activity.

### Chemical screens identify novel AHR inducers

AHR is activated by a diverse array of chemicals, including microbial and dietary indole derivatives, endogenous tryptophan metabolites, polycyclic aromatic hydrocarbons, and flavonoids (1, 54-56). Prior characterization of regulators of AHR signaling had suggested that regulators vary depending on the identity of the inducer, with some regulators being specific to individual AHR inducers (38, 40-42). We next sought to ask whether the regulators of AHR signaling identified in our genome-scale screens are shared across different AHR inducers. For this purpose, we first aimed to compile a set of chemicals from different classes that activate AHR in HepG2 cells, by evaluating previously reported AHR inducers and screening for potential new inducers.

To enable high-throughput screening of chemicals for AHR activation, we modified our GFP-based AHR reporter into a dual luciferase reporter (Figure 2A). This reporter comprises AREs upstream of a minimal promoter (p(cFos) or p(TATA) containing a TATA-box promoter element) that drives expression of firefly luciferase. To enable signal normalization and isolation of transduced cells, the reporter also contains a renilla luciferase-P2A-GFP cassette driven by a constitutive promoter (UBC). A polyclonal HepG2 cell line stably transduced with the reporter responded to I3S with high dynamic range and with a dose-response curve that matched induction of a canonical AHR target gene, *CYP1A1* (Figure 2B). 13 additional known AHR inducers also activated the reporter to different extents, including weak but clearly detectable responses to compounds reported to be relatively weak AHR activators, such as cinnabarinic acid (57) (Figure S2A). Thus, our luciferase reporter has the dynamic range to identify AHR activators across a range of potencies.

**Figure 2.**
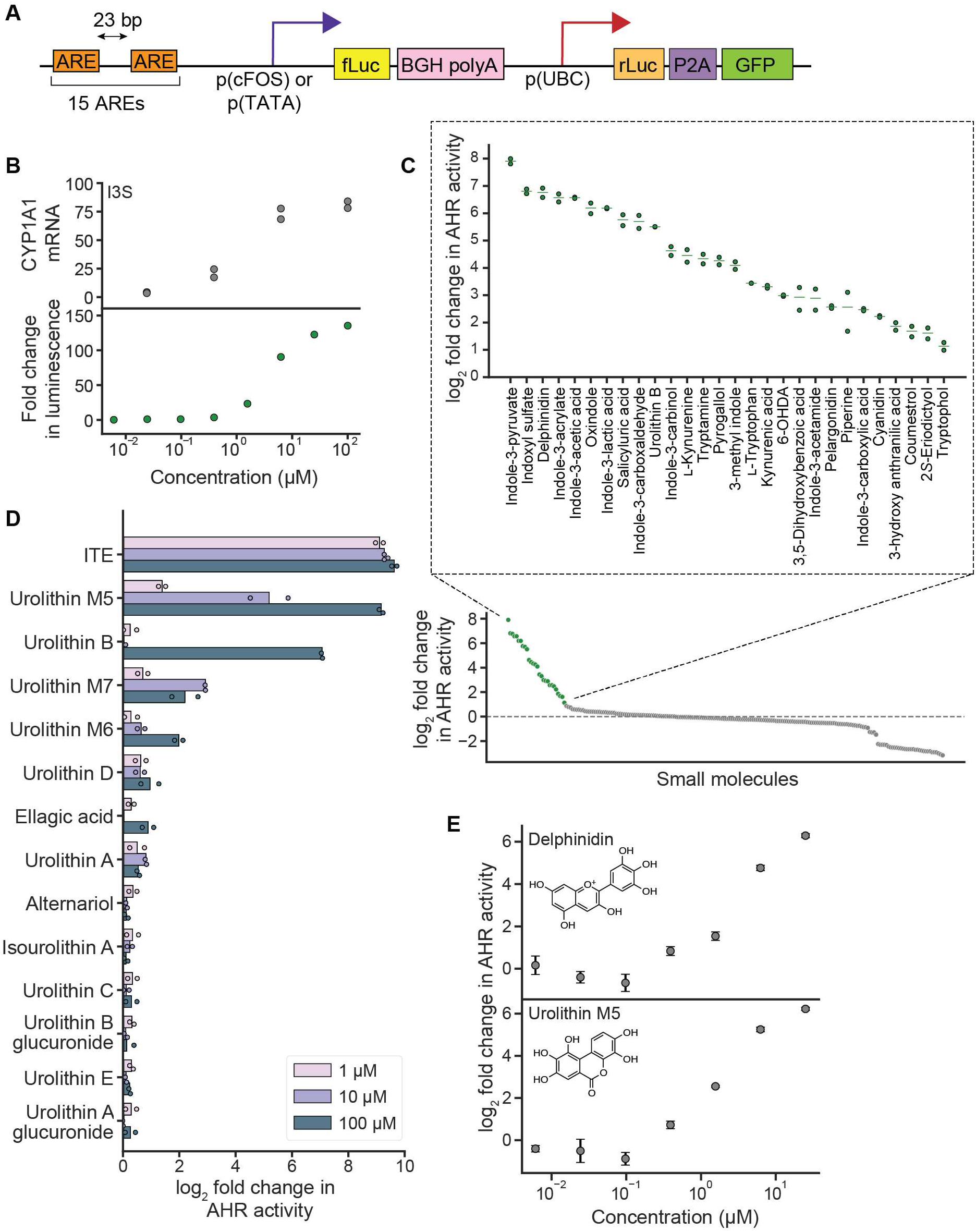
Chemical screens identify novel AHR inducers. (A) Schematic of luciferase-based AHR reporter. (B) Induction of *CYP1A1* expression (top, measured by RT-qPCR) and induction of the luciferase reporter (bottom) after treatment with I3S for 4 h. (C) log_2_ fold change in AHR luciferase reporter signal after treatment with the microbiome-derived small molecule library for 4 h, normalized to corresponding solvent controls. Dashed line indicates log_2_ fold change = 0. Inset shows all molecules with log_2_ fold change > 1. (D) log_2_ fold change in AHR luciferase reporter signal by urolithin and related compounds, normalized to solvent control. ITE is a positive control. (E) Dose-response curves for the strongest novel inducers from each screen, delphinidin and urolithin M5, after treatment of luciferase reporter cells for 4 h. Data represent mean ± standard deviation (*n*=3).

Next, we conducted chemical screens using two sets of chemicals: (1) a library of ∼240 small molecules associated with the human microbiome that we curated (Figure 2C, Table S3) and (2) 13 urolithins and related compounds, which are derived from gut microbial metabolism of the polyphenol ellagic acid and have been implicated as activators of AHR (Figure 2D, Table S4) (26, 58). For each screen, we treated the HepG2 reporter cells with each molecule for 4 h, measured the ratio of firefly luciferase and renilla luciferase signal, and then calculated the log_2_ fold change relative to corresponding vehicle controls. Across the two sets of chemicals, we identified 32 compounds that were not toxic and induced AHR with a log_2_ fold change ≥ 1 (Figure 2C-D, S2B). These compounds included many previously known AHR inducers from different chemical classes, including multiple indole derivatives (11, 24, 55), kynurenine (15, 59), and the flavonoid pelargonidin (60) (Figure 2C). Of the previously reported AHR inducers in our chemical libraries, only urolithin A did not affect AHR activity at any tested concentration (Figure 2D), but urolithin A has been disparately described as an AHR agonist as well as an antagonist (26, 58,61, 62). Our work adds to this body of literature to suggest that the role of urolithin A in AHR activation depends on context.

Finally, the AHR inducers in our chemical screens also included compounds not previously known to activate AHR, such as delphinidin, salicyluric acid, pyrogallol, and urolithin M5 (Figure 2C-D). Further examination of dose-response relationships for these compounds revealed that delphinidin and urolithin M5 robustly induce AHR signaling at low micromolar concentrations (Figure 2E, S2C-D). Both compounds are produced by gut bacterial metabolism of dietary polyphenols, which have been estimated to reach µM to mM concentrations in the intestine, depending on dietary intake (63). Thus, these compounds may contribute to AHR activity in vivo. In the absence of binding assays, we cannot formally establish whether these compounds induce AHR signaling by acting as AHR ligands or by indirect mechanisms. For this reason, we refer to these compounds as inducers of AHR signaling throughout the remainder of the manuscript. In sum, our chemical screens recapitulated known patterns of AHR activation and identified potentially novel inducers of AHR signaling.

### Sublibrary CRISPRi screens reveal AHR regulatory networks across inducers

With this collection of AHR inducers, we sought to ask whether the AHR regulators identified in our genetic screens play similar roles in activation of AHR by different inducers. For this purpose, we first constructed a sublibrary of sgRNAs targeting genes for which knockdown had strong effects on AHR activation by either I3S or IPY in the genome-scale screen. We excluded several genes in essential processes, such as many ribosomal and proteasomal genes, because we reasoned that knockdown of such genes likely influenced AHR activity indirectly. We also included 66 genes from categories that had been linked to AHR activation by other inducers, including transporters, tryptophan metabolism enzymes, and transcriptional coregulators. The final library comprised 2715 sgRNAs targeting 497 genes (543 TSSs with 5 sgRNAs per TSS) and 150 non-targeting sgRNAs.

With this sublibrary, we conducted FACS-based pooled screens to identify modulators of AHR activation by 14 different inducers (12 known inducers, delphinidin, and urolithin M5; Figure 3A-B, S3A, Table S5-S7), in the same manner as the genome-scale screen. We also conducted a growth screen to define how knockdown of these genes affects growth of HepG2 cells (Figure 3B, Table S8-S10). For this purpose, we harvested a population of sgRNA-expressing cells at the outset of the experiment, cultured a second population of cells for 10 doublings, and quantified the differences in sgRNA abundances between those two timepoints. Phenotypes for both AHR activation and growth were well-correlated between replicates (Figure S3B-C). Further, gene-level AHR activation phenotypes were well-correlated between the sublibrary and genome-wide screens for I3S and IPY (*r*^2^ = 0.66 and 0.71 for I3S and IPY, respectively; Figure 3C), underscoring the highly reproducible nature of our screens. Using data from the growth screen, we excluded 157 genes for which knockdown caused strong growth defects (growth phenotype < −0.45, Figure 3D, S3D) from further analysis because AHR activation phenotypes for these genes were more variable due to low sgRNA counts at the end point and because knockdown of these genes likely affects AHR signaling indirectly.

**Figure 3.**
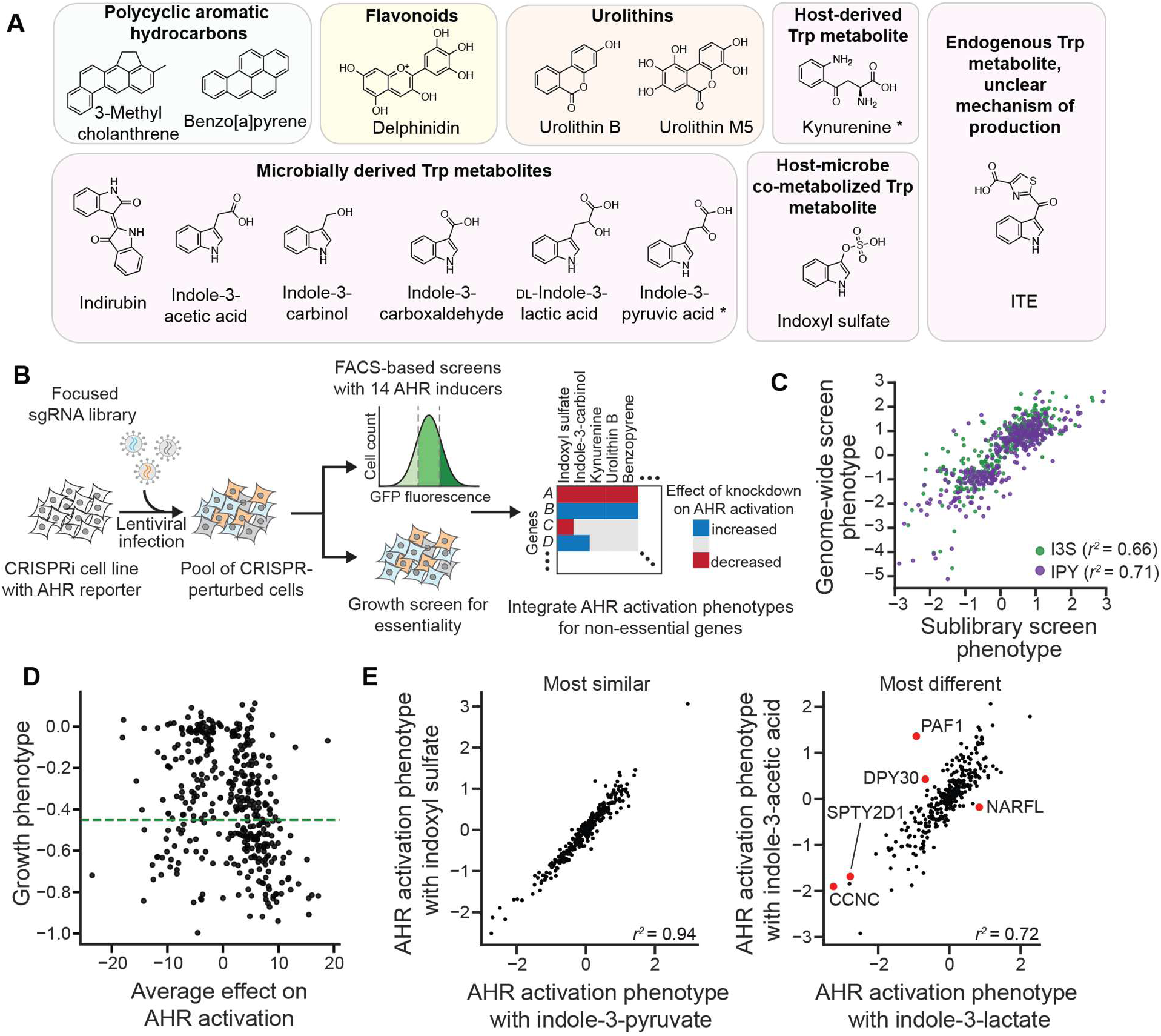
Sublibrary CRISPRi screens reveal AHR regulatory networks across inducers. (A) Chemical structures of small molecules used in the sublibrary CRISPRi screens grouped by chemical class. Kynurenine and indole-3-pyruvate (marked with *) may be independently derived from both host and microbial sources. (B) Schematic of FACS-based and growth-based sublibrary CRISPRi screens. (C) Correlation between AHR activation phenotypes in the genome-wide and sublibrary screens for I3S and IPY. (D) Growth phenotypes for all genes that are hits with at least 1 inducer plotted against their average z-scored AHR activation phenotypes across inducers. Dashed line shows the threshold used to classify genes as essential. (E) Comparison of AHR activation phenotypes of the two most similar (left) and two most different (right) inducers based on the correlation between the AHR activation phenotypes of all genes that are hits with at least one inducer. The 5 genes whose phenotypes are most different between the two inducers (right) are shown in red. *r*^2^ is squared Pearson correlation coefficient.

Profiles of AHR activation phenotypes are well-correlated across most inducers (Figure S3E). These profiles are nearly identical for some inducers, such as for I3S and IPY, but have some differences for others, such as for indole-3-lactate and indole-3-acetic acid (Figure 3E). Chemical similarity alone does not predict these differences. For example, the phenotype profile of 3-methylcholanthrene is more similar to that of indirubin than to that of the more chemically similar compound benzo[a]pyrene. Thus, the regulatory networks of different inducers are composed of some factors that are shared and some that are inducer-specific.

### Identifying general and inducer-specific regulators of AHR activity

We next asked which regulators are shared or different between inducers. Some regulators affected AHR activation by all inducers, whereas others only affected activation by a subset (Figure 4A, Table S11). Grouping genes by AHR activation phenotypes revealed three main categories: (i) genes for which knockdown affects AHR activation in the same direction with all inducers (general regulators), (ii) genes for which knockdown affects AHR activation in opposite directions with different inducers (divergent regulators), and (iii) genes for which knockdown affects AHR activation by a subset of inducers but not by others (inducer-specific regulators) (Figure 4B-E). Each category expands our understanding of the regulation of AHR signaling, as we outline next.

**Figure 4.**
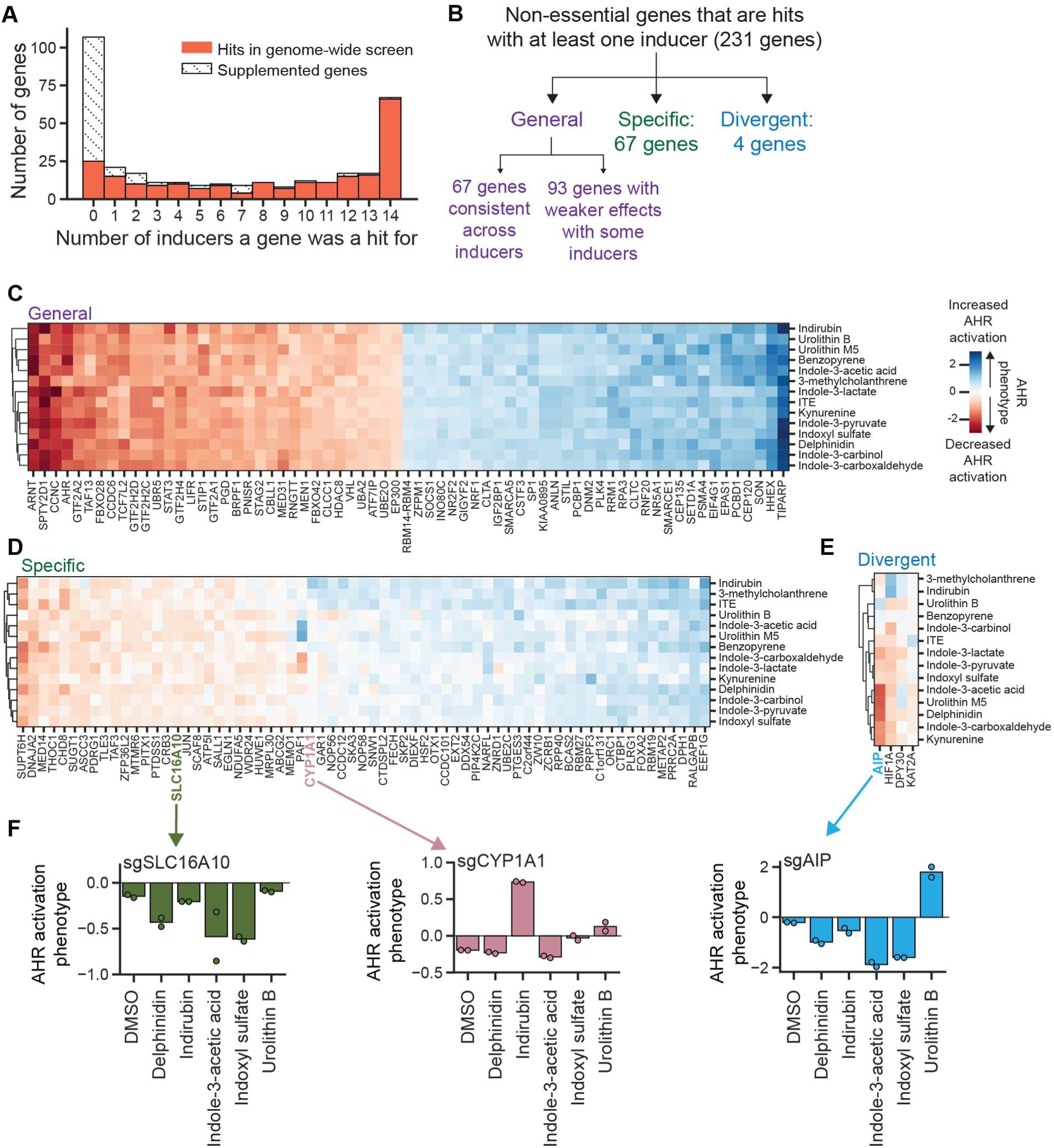
Identification of general and inducer-specific regulators of AHR signaling. (A) Distribution of the number of inducers for which a gene was a hit, separated by whether they were hits in the genome-wide screen or supplemented. (B) Classification scheme for regulators of AHR activity. (C) AHR activation phenotypes for all genes classified as general regulators. (D) AHR activation phenotypes for all genes classified as specific regulators. PAF1 has divergent AHR activation phenotypes between inducers but has low *p* values, likely due to a strong growth phenotype (-0.42) resulting in low counts. It therefore did not pass the threshold for classification as a divergent regulator. (E) AHR activation phenotypes for all genes classified as divergent regulators. (F) AHR activation phenotypes of two specific (SLC16A10, CYP1A1) and one divergent regulator (AIP) from independent validation, measured by transducing individual sgRNAs and measuring GFP fluorescence by flow cytometry after treatment with indicated inducers for 24 h (delphinidin, indole-3-acetic acid, indoxyl sulfate, urolithin B: 100 µM; indirubin: 10 µM).

67 genes classified as general regulators of AHR activation by our most stringent threshold for calling hits (Figure 4C). These genes likely act through central nodes of the AHR signaling pathway, such as expression of AHR, nuclear translocation, interaction with coregulators, or DNA binding. These genes include known regulators of AHR signaling, including *ARNT* and *TIPARP*, as well as many genes not previously known to regulate AHR signaling, including components of the ubiquitin modification system (*UBR5, RNF20, UBA2, UBE2O*), epigenetic modifiers (*HDAC8, SMARCA5, SMARCE1, SETD1A*), and components of the JAK1-STAT3 signaling pathway (*LIFR, STAT3*, and *SOCS1*). Knockdown of 93 additional genes affected AHR activation by all inducers but with weaker effects for some inducers (Figure S4A). These genes include additional ubiquitinating enzymes (*UBQLN4*), epigenetic modifiers (*SETDB1, INO80C*), and components of the JAK1-STAT3 signaling pathway (*JAK1*).

Another 67 genes classified as inducer-specific regulators of AHR activation (Figure 4D). These genes have different patterns of specificity: knockdown of some genes affects AHR activation by all but one inducer, knockdown of other genes affects AHR activation only by a single inducer, and knockdown of yet other genes affects AHR activation by a subset of inducers. For example, knockdown of *SUPT6H*, which encodes a transcriptional elongation factor, reduces AHR activation by all inducers except urolithin B; knockdown of *ABCG2*, which encodes an efflux pump and was not a hit in the genome-scale screen but which we had supplemented into the sgRNA sublibrary, reduces AHR activation by urolithin M5 but does not affect activation by other inducers; and knockdown of *SLC16A10*, which encodes an aromatic amino acid transporter, markedly reduced AHR activation for certain inducers such as delphinidin and indole-3-acetic acid but had little to no effect on others such as 3-methylcholanthrene (Figure 4D, 4F). We recapitulated these observations using individual knockdowns (Figure 4F). The inducer-specific regulators also included known regulators of AHR signaling, such as *CYP1A1*, for which knockdown increased AHR activation in the presence of 3 inducers: indirubin, 3-methylcholanthrene, and, to a lesser extent, benzo[a]pyrene (Figure 4D, 4F). *CYP1A1* encodes a cytochrome P450 enzyme that negatively regulates AHR activity by metabolizing and thereby inactivating some AHR inducers (32, 64, 65). This activity is best characterized for the strong AHR inducer 6-formylindolo[3,2-*b*]carbazole (FICZ). Our observations suggest that CYP1A1 also metabolizes these three inducers but not the others in our set, corroborating and extending previous findings on substrate preferences of CYP1A1 (66-69).

Finally, 4 genes, *AIP, HIF1A, DPY30,* and *KAT2A*, classified as divergent regulators of AHR activation (Figure 4E). Perhaps the most surprising gene in this category is AIP (AHR interacting protein, also known as *XAP2* or *ARA9*), which binds to AHR in the cytosol as part of an HSP90 chaperone complex and is thought to be generally required for AHR activity (70). Knockdown of *AIP* resulted in the expected decrease in AHR activation for most inducers but caused an increase in AHR activity in the presence of urolithin B (Figure 4E). We confirmed this observation using individual knockdown of *AIP* in our GFP and luciferase reporter cell lines (Figure 4F, S4B). One possible explanation for this observation is that most inducers require AHR to be complexed with AIP, whereas urolithin B binds to AHR that is not complexed with AIP. Because unliganded AHR is thought to be degraded in the absence of AIP, a prediction of this model is that in the absence of AIP, addition of urolithin B, but not other inducers, stabilizes AHR. Quantification of AHR protein levels in cells with knockdown of *AIP*, however, revealed no difference in AHR levels in cells treated with urolithin B or indole-3-acetic acid (Figure S4C), suggesting that the divergent role of AIP is not explained by this model. We expect that direct binding or structural assays will be required to define this mechanism.

In sum, we identify regulators that are shared across all inducers, regulators that only contribute to AHR activation by some inducers, and regulators that have divergent roles in AHR activation by different inducers.

### Functional classification of general regulators of AHR signaling

Our screens collectively nominated many previously unappreciated genes as general regulators of AHR activity. We reasoned that further defining mechanisms of action for these regulators would deepen our understanding of the core regulatory network that underpins AHR activation. We selected 43 genes for which knockdown modulated AHR signaling by all tested inducers (37 general according to our most stringent threshold and 6 likely general that had weaker effects with a subset of inducers) and used a stepwise approach to pinpoint at which node of the AHR signaling pathway each gene acts.

First, we asked whether knockdown of these genes affects mRNA levels of AHR at baseline. Knockdown of 6 genes strongly affected *AHR* mRNA levels (absolute log_2_ fold change relative to non-targeting control sgRNAs ≥ 1) in the same direction as AHR activation (Figure 5A). These genes include transcription factors (*HHEX, TCF7L2*), chromatin-modifying enzymes (*SMARCE1, RNF20*) and ubiquitin ligases (*FBXO42*), suggesting that varied cellular process contribute to regulation of *AHR* mRNA levels.

**Figure 5.**
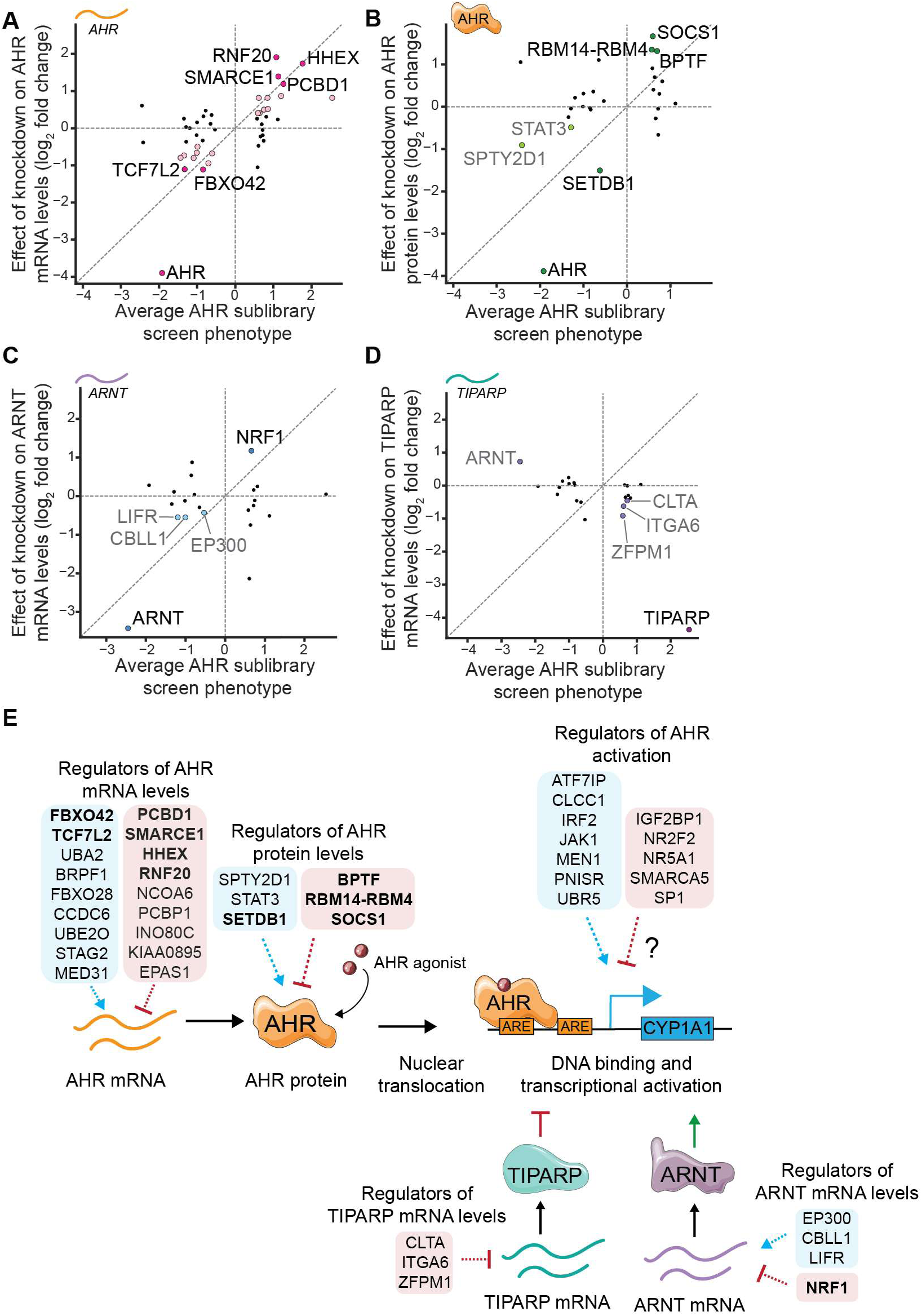
Functional classification of general regulators of AHR signaling. Effect of knockdown of selected general regulators on (A) AHR mRNA levels, (B) AHR protein levels, (C) ARNT mRNA levels, and (D) TIPARP mRNA levels, plotted against their average AHR sublibrary screen phenotype across inducers. Genes with strong effects (log_2_ fold change > 1) are highlighted in dark colors and genes with moderate effects (log_2_ fold change between 0.4 and 1) are highlighted in lighter colors. Knockdown of ZFPM1, NR2F2, ITGA6 and SP1 result in log_2_ fold change > 0.4 for AHR protein levels but are not classified due to inconsistent effects across replicates (Figure S5B). (E) Summary of identified roles of general regulators of AHR signaling. Bolded genes are the ones with strong effects (log_2_ fold change > 1).

Knockdown of 12 additional genes led to moderate changes in *AHR* mRNA levels (absolute log_2_ fold change relative to non-targeting control sgRNAs between 0.4 and 1). These genes include additional post-translational modification enzymes (*UBA2, UBE2O, FBXO28*) as well as chromatin-modifying enzymes (*BRPF1, INO80C*).

One of the strongest regulators of *AHR* mRNA levels is HHEX, a homeobox transcription factor that had not previously been linked to AHR expression. We find that loss of HHEX leads to increased AHR activation in our screens and increased *AHR* transcript levels (Figure 5A, S5A). Subsequent western blotting confirmed that knockdown of *HHEX* also leads to increased AHR protein levels (Figure S5B). Several independent lines of evidence further corroborate our results. First, in an RNA-seq dataset of early pancreatic progenitor cells, HHEX knockout cells have increased AHR mRNA levels compared to wild-type cells (71). Second, ChIP-seq data for HHEX contain peaks upstream of the *AHR* gene in both ESC-derived pancreatic cells (71, Figure S5C) and HCT-116 colon cancer cells (72). Finally, examining the relationship between transcript levels of *HHEX* and *AHR* across cell lines using data from the Human Protein Atlas revealed that cells with high *HHEX* transcript levels invariably have low *AHR* transcript levels (Figure S5D). Our results together with these observations suggest that HHEX binds at the *AHR* locus and inhibits expression of *AHR*.

Knockdown of 25 genes did not appreciably affect *AHR* mRNA levels. We next measured how knockdown of these genes affects AHR protein levels using western blotting. Knockdown of 4 genes led to large changes in AHR protein levels (absolute log_2_ fold change relative to non-targeting control sgRNAs ≥ 1) and knockdown of 2 genes led to moderate changes (absolute log_2_ fold change relative to non-targeting control sgRNAs between 0.4 and 1, Figure 5B, S5B) in the same direction as AHR activation phenotypes. Several of these genes are chromatin modifiers (*BPTF, SETDB1, SPTY2D1*), knockdown of which may influence AHR protein levels indirectly. We were most intrigued by changes to AHR protein levels induced by knockdown of *STAT3* and *SOCS1*, which are components of the JAK-STAT signaling pathway.

Specifically, we find that knockdown of *STAT3* decreases AHR protein levels and knockdown of *SOCS1*, a negative regulator of STAT3, increases AHR protein levels, implying that STAT3 signaling stabilizes AHR at the protein level at baseline. Indeed, chemical inhibition of STAT3 signaling leads to acute depletion of AHR protein (Figure S5E). Our work complements previous observations that oncostatin M-induced STAT3 signaling upregulates AHR expression (73) and that AHR can induce STAT3 expression (74), further emphasizing bi-directional crosstalk between these two signaling pathways.

Finally, we measured how knockdown of the remaining 19 candidate general regulators affects expression of two core regulators of AHR, ARNT and TIPARP (Figure S5F). Knockdown of 1 gene strongly and knockdown of 3 genes moderately affected *ARNT* mRNA levels (Figure 5C), and knockdown of 3 genes moderately affected *TIPARP* mRNA levels, as assessed using the thresholds described above (Figure 5D). We observed the largest effect upon knockdown of nuclear respiratory factor 1 (NRF1), a transcription factor with broad roles in regulation of metabolism and mitochondrial function, for which knockdown led to a 2.3-fold increase in *ARNT* mRNA levels (Figure 5C). ChIP-seq tracks from the ENCODE database indicate putative NRF1 binding sites upstream of the *ARNT* promoter (Figure S5G). Thus, NRF1 may inhibit AHR activation by inhibiting ARNT expression.

In summary, our tiered approach revealed general regulators that act at different stages of the AHR signaling pathway, including modulators of *AHR* mRNA and protein levels and modulators of expression of the core regulators ARNT and TIPARP (Figure 5E). Knockdown of 12 genes did not influence AHR signaling at the stages that we assayed; these genes are candidate regulators of AHR activation and may for example act at the level of post-translational modification of AHR, nuclear translocation, DNA binding, coactivator recruitment, or turnover (Figure 5E).

### Many general regulators of AHR signaling act in a cell type-specific manner

AHR is expressed in most cell types in humans, and some factors regulate AHR signaling in a cell type-specific manner (43, 75). We therefore asked if genes we had found to be general regulators of AHR activation in HepG2 cells are also involved in regulation of AHR activation in U-87 glial cells. AHR has been linked to oncogenesis in glioma, making U-87 cells a relevant model to interrogate the genetic networks that mediate AHR induction (76). We confirmed using our luciferase reporter system that AHR inducers strongly activate AHR in U-87 cells, albeit less strongly than in HepG2 cells (Figure S6A). We generated U-87 cells with stable expression of Zim3-dCas9, confirmed that these cells have strong knockdown of endogenous genes (Figure S6B), integrated our AHR GFP reporter, and isolated a monoclonal cell line. The resulting AHR reporter U-87 cells display a homogenous increase in GFP signal after treatment with 100 µM I3S or 100 µM IPY for 24 h, as measured by flow cytometry (Figure 6A). CRISPRi-mediated knockdown of AHR abolished reporter induction by 100 µM IPY whereas knockdown of TIPARP amplified reporter induction, confirming that the response is specific to AHR signaling (Figure 6B).

**Figure 6.**
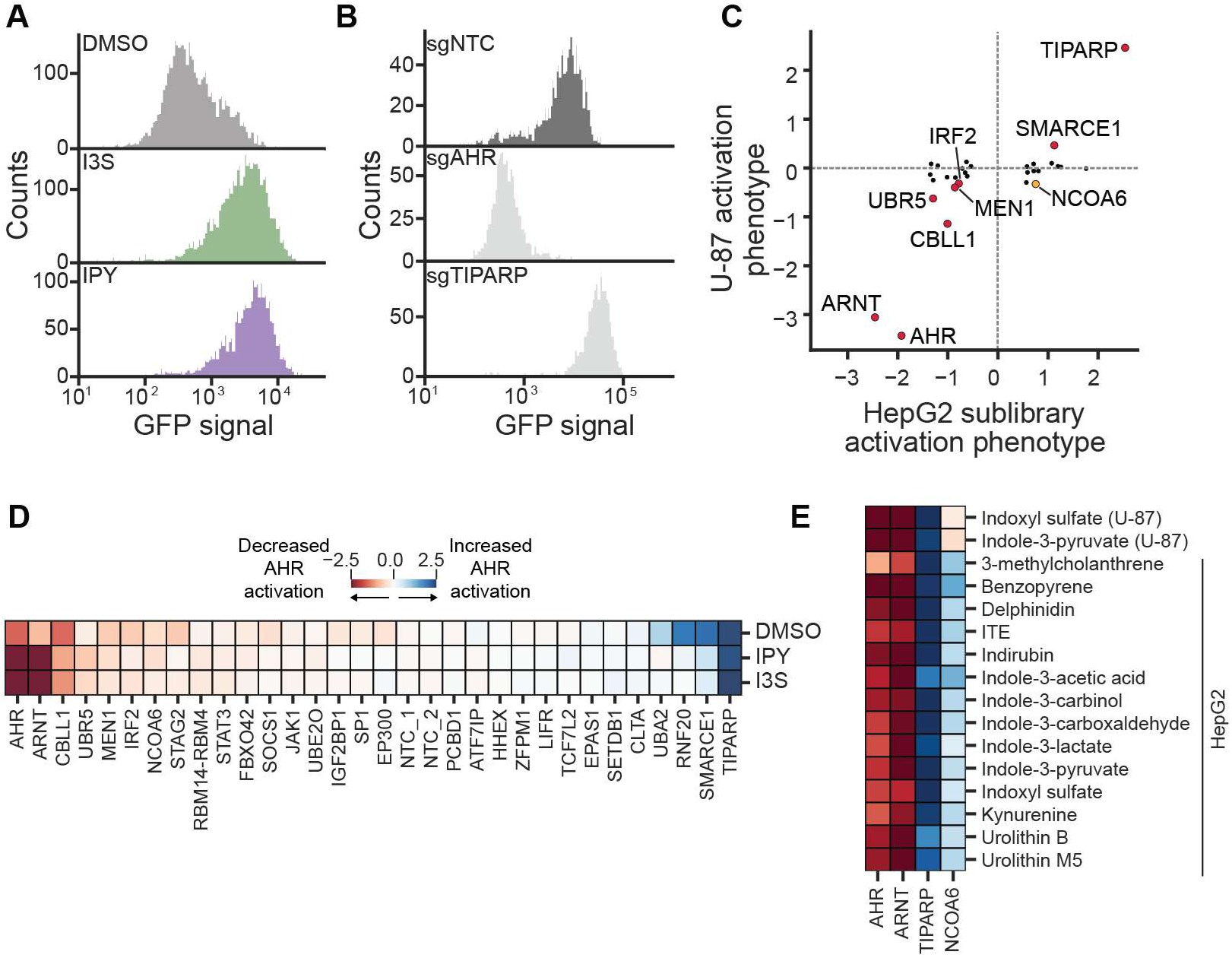
Many general regulators of AHR signaling act in a cell type-specific manner. (A) GFP fluorescence of monoclonal U-87 reporter (cKD004) cells treated with 100 µM I3S or 100 µM IPY for 24 h, as measured using flow cytometry. (B) GFP fluorescence of U-87 reporter cells transduced with the indicated sgRNAs and treated with 100 µM IPY for 24 h. Data are representative of two independent transductions. sgNTC is a non-targeting control sgRNA. (C) Average AHR activation phenotypes of genes in U-87 cells plotted against their average HepG2 sublibrary screen phenotypes. Red points indicate genes that have similar effects in both cell types. The yellow point indicates a gene with opposite phenotypes in the two cell types. (D) AHR activation phenotypes in U-87 cells for all genes with 100 µM I3S, 100 µM IPY, or vehicle control. (F) AHR activation phenotypes in U-87 and HepG2 cells for 3 core regulators of AHR signaling (AHR, ARNT, TIPARP) and one gene categorized as a general regulator in HepG2 cells (NCOA6).

We next measured how knockdown of the 43 general regulators of AHR activation that we had characterized in HepG2 cells affected AHR activation by I3S and IPY in U-87 cells using our arrayed, internally controlled assays (Figure S6C). We excluded from further analysis 16 genes for which < 100 sgRNA-expressing cells remained at the endpoint. Among the remaining 27 genes, knockdown of 5 genes, *CBL11*, *IRF2, MEN1*, *SMARCE1*, and *UBR5*, led to a change in AHR activation that was of similar magnitude and direction as in HepG2 cells (absolute activation phenotypes ≥ 0.3) (Figure 6C). We noticed that in U-87 cells, unlike in HepG2 cells, knockdown of several genes, including *AHR*, *ARNT*, and *TIPARP*, led to changes in reporter signal at baseline. For some genes, including *SMARCE1*, *RNF20*, and *STAG2*, the effect of knockdown on AHR activity is stronger at baseline than with inducer treatments (Figure 6D). These observations point to higher basal AHR signaling in U-87 cells relative to HepG2 cells and may indicate that RNF20 and STAG2 regulate AHR similarly in both cell types.

Knockdown of most of the remaining genes did not strongly affect AHR activity in U-87 cells (Figure 6D, S6D). 6 of these genes are not expressed in U-87 cells, as determined using expression data from the Human Protein Atlas, providing an explanation for the lack of effect on AHR activation (Figure S6E). The lack of effect for the remaining genes is not explained by a lack of expression or by lack of knockdown (Figure S6F), suggesting that these factors do not contribute to regulation of AHR activation in U-87 cells.

Perhaps most surprisingly, knockdown of the transcriptional coregulator NCOA6 led to opposite effects on AHR activation in the two tested cell types: an increase in AHR activation in HepG2 cells and a decrease in AHR activation in U-87 cells (Figure 6E). Previous work has implicated differential recruitment of transcriptional coregulators in mediating differential activities of AHR in different cell types (43). The opposite effect of knockdown of NCOA6 on AHR signaling in HepG2 and U-87 cells may point to such a mechanism.

Thus, although some factors contribute to regulation of AHR activation across cell types, many other factors are specific to individual cell types, including some that exert opposite effects in different cell types.

### The E3 ubiquitin ligase UBR5 is a regulator of AHR turnover

Finally, we were particularly interested in further characterizing general regulators that regulated AHR activity in both cell types and whose effect was not explained by effects on *AHR* mRNA or protein levels. We focused on UBR5, a HECT-family E3 ubiquitin ligase that has garnered substantial attention for its roles in degradation of unpaired subunits of multi-subunit transcription regulators (orphan quality control) (77), degradation of agonist-bound nuclear receptors (78), and regulation of MYC levels (79). In our experiments, knockdown of *UBR5* reduced AHR activity by all tested inducers and in both cell types (Figure 7A). Further, knockdown of *UBR5* did not affect mRNA levels of *AHR*, *ARNT*, or *TIPARP* or protein levels of AHR at baseline (Figure 5A-D, S7A), suggesting that UBR5 contributes to regulation of AHR activation via an unknown mechanism.

**Figure 7.**
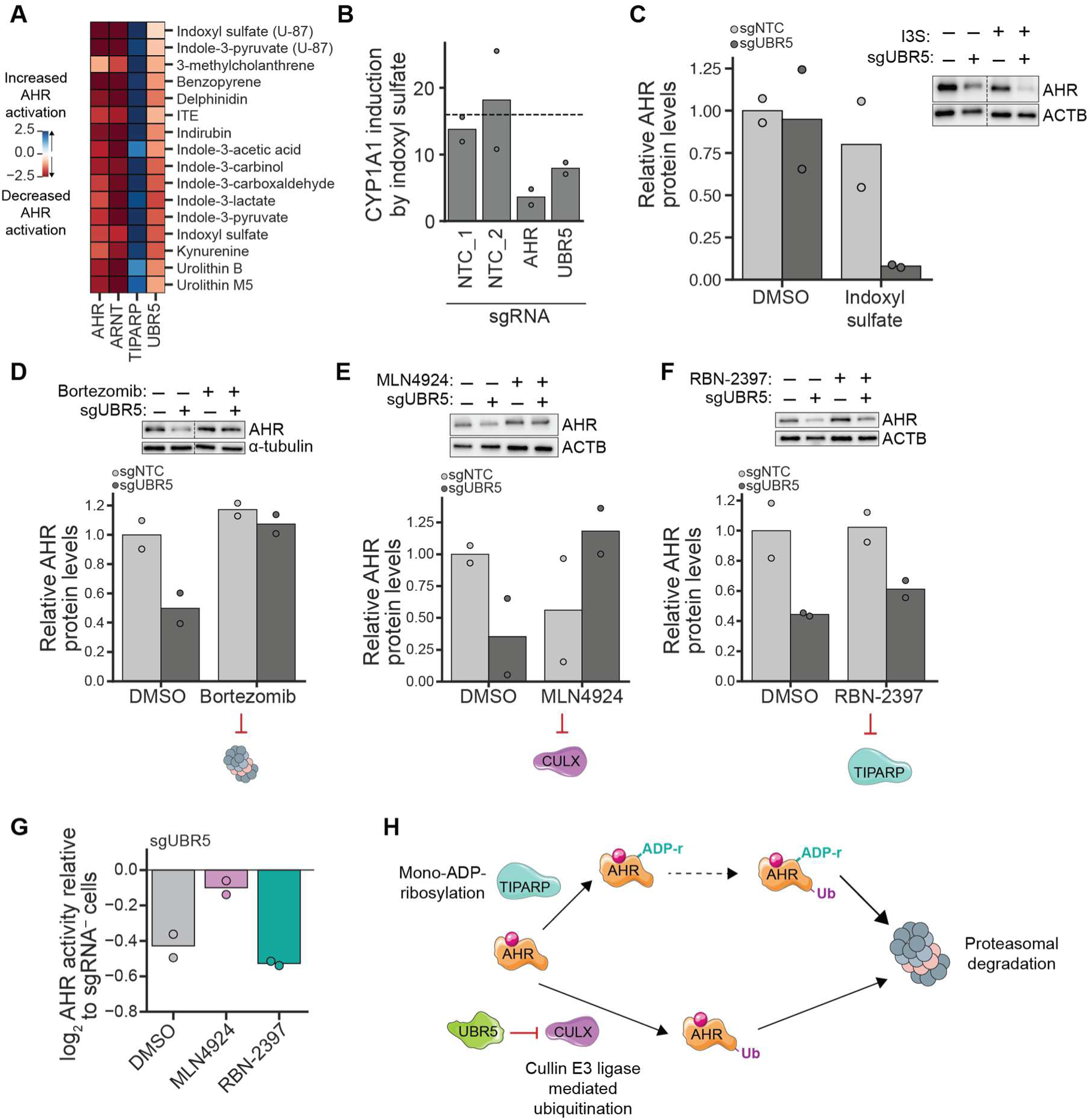
The E3 ubiquitin ligase UBR5 is a regulator of AHR turnover. (A) AHR activation phenotypes in U-87 and HepG2 cells for 3 core regulators of AHR signaling (AHR, ARNT, TIPARP) and UBR5. (B) *CYP1A1* mRNA levels in HepG2 cells with the indicated knockdowns after treatment with 10 µM I3S for 4 h relative to vehicle-treated cells, measured using RT-qPCR. Dashed line indicates the average induction in the two non-targeting control (NTC) cell populations. (C) Left: Relative AHR protein levels in HepG2 cells with either a non-targeting control sgRNA (NTC) or UBR5 knockdown treated with 100 µM I3S for 4 h, measured by Western blotting. Right: representative blot. The dashed line indicates wells run non-contiguously. Additional images in Figure S7B. (D) Bottom: Relative AHR protein levels in HepG2 cells with either a NTC or UBR5 knockdown pre-treated with 500 nM bortezomib or vehicle for 2 h and treated with 100 µM IPY for 4 h. Top: representative blot. The dashed line indicates wells run non-contiguously. Additional images in Figure S7C. (E) Bottom: Relative AHR protein levels in HepG2 cells with either a NTC or UBR5 knockdown pre-treated with 1 µM MLN4924 or vehicle for 2 h and treated with 100 µM I3S for 4 h. Bottom: representative blot. Additional images in Figure S7D. (F) Bottom: Relative AHR protein levels in HepG2 cells with either a NTC or UBR5 knockdown pre-treated with 1 µM RBN-2397 or vehicle for 4 h and treated with 100 µM I3S for 4 h. Bottom: Representative blot. Additional images in Figure S7E. (G) AHR activation phenotypes in UBR5 knockdown cells pre-treated with either 1 µM MLN4924 or 1 µM RBN-2397 for 2 h before treatment with 100 µM IPY for 4 h. AHR activation phenotypes were calculated using the log_2_ ratio of median GFP fluorescence in sgRNA-expressing cells relative to median GFP fluorescence for non-sgRNA-expressing cells in the same well. (H) Proposed model for regulation of AHR protein levels by UBR5.

We first confirmed that knockdown of *UBR5* reduces induction of *CYP1A1* by I3S in HepG2 cells (Figure 7B). To further investigate the role of UBR5 in AHR signaling, we asked whether UBR5 influences AHR levels in the presence of an inducer. Strikingly, cells with knockdown of *UBR5* had ∼10-fold lower AHR protein levels than control cells after treatment with I3S for 4 h (Figure 7C, S7B). We thus hypothesized that UBR5 plays a role in regulating AHR protein levels after induction.

Thus far, two factors have been implicated in AHR degradation (80): TIPARP and the cullin CUL4B. TIPARP mediates mono-ADP-ribosylation of AHR, likely followed by ubiquitination (33, 34), although the specific E3 ligases that ubiquitinate AHR remain unidentified. It also remains unclear whether TIPARP and CUL4B act in a linear pathway or independently to regulate AHR degradation. We used chemical inhibitors to define if AHR is degraded by one of these pathways in the absence of UBR5. Inhibition of the proteasome using bortezomib and inhibition of cullin-containing E3 ligases using a neddylation inhibitor (MLN4924) (81) prevented the decrease in AHR protein levels upon treatment with an AHR agonist in cells with knockdown of *UBR5* (Figure 7D-E, S7C-D). By contrast, treatment with a TIPARP inhibitor (RBN-2937) (82) did not prevent loss of AHR (Figure 7F, S7E). We confirmed these findings using internally normalized reporter assays (Figure 7G): inhibition of neddylation but not inhibition of TIPARP reverted the decrease in AHR activation in cells with knockdown of *UBR5*. TIPARP inhibition using RBN-2397 increased AHR activation in control cells (Figure S7F), confirming that the inhibitor was functional.

Together, these findings support a model in which loss of UBR5 leads to ubiquitination of AHR by a cullin RING ligase (CRL) complex and subsequent degradation by the proteasome, in a manner that is independent of TIPARP (Figure 7H). Because no cullins were hits in our screens but CUL4B has previously been reported to promote ubiquitination of AHR, the leading candidate is a CRL that can use either CUL4A or CUL4B. Finally, it remains to be determined if UBR5 restrains the activity of this degradation pathway at baseline, for example by ubiquitinating and mediating degradation of a component of a CRL complex, or if this pathway is activated in an indirect manner in the absence of UBR5. Our results reveal a novel role for UBR5 in regulation of AHR signaling and point to a distinct pathway by which levels of ligand-bound AHR are regulated.

## Discussion

Several decades of research have established multifaceted physiological roles for AHR signaling, including in regulation of xenobiotic detoxification, intestinal barrier integrity, and T cell polarization. Both loss and overactivation of AHR have now been implicated in the etiology of metabolic and inflammatory diseases. Many of these insights, however, have been derived from experimental models with complete knockout or massive activation of AHR, leaving open the identities of factors that contribute to regulation or dysregulation of AHR activity in physiological settings. Our genome-scale genetic screens now establish a comprehensive set of factors whose levels contribute to regulation of AHR activity in a model of human hepatocytes. Many are shared across all AHR inducers that we tested and thus are likely to be general regulators in this cell type. A further strength of our experimental system is that it reports on the transcriptional activity of AHR with high dynamic range, using cognate binding motifs, and at endogenous expression levels of AHR, which allows us to identify factors along the entire AHR signaling pathway. By further characterizing a subset of these factors, we place them at different steps of the pathway. For example, we find that HHEX regulates transcription of *AHR*. We also identify factors that affect AHR activity without changing its expression. For example, the E3 ubiquitin ligase UBR5 promotes stability of ligand-bound AHR (Figure 5E). The hypotheses derived from this work as well as the databases of identified regulators, which are included in full as supplementary data (Tables S1-S3, S6-11), establish a resource for further interrogation of mechanisms of AHR signaling.

Many of the regulators we discovered fulfil additional roles beyond their involvement in AHR signaling. For example, UBR5 also mediates degradation of unpaired transcription factor subunits (77), DNA-bound nuclear receptors (78), and MYC (79). NRF1, which we propose suppresses AHR signaling by repressing *ARNT* expression, broadly regulates expression of metabolic and mitochondrial genes. These findings reflect a common theme in AHR signaling. Indeed, even ARNT, a core component of the AHR signaling pathway, is known to fulfil additional roles, including as a dimerization partner for HIF-1α. Our work thus underscores how integrated AHR signaling is into the signaling circuits of human cells.

Another intriguing aspect of AHR signaling is the reported variability and context-dependence. Some instances of context-dependent signaling result from post-translational modifications of AHR and recruitment of specific coregulators (13, 40, 41), but in many cases the molecular bases have remained unclear. We find that many regulators are specific to individual cell types or inducers, and some have opposite effects on AHR activation in different cell types, such as NCOA6, or by different inducers, such as AIP. Even some factors in the core AHR signaling pathway have context-specific roles. In addition to AIP, AHRR did not score in our screens, likely because it is not expressed in HepG2 cells, and CYP1A1 scored as a negative regulator of AHR signaling only for specific inducers, consistent with previous reports (66-69). A limitation of our work in this regard is that the two inducers we chose for our initial genome-scale screens, indoxyl sulfate and indole-3-pyruvate, turned out to have the most similar phenotype profiles among inducers we tested in our sublibrary screens. Thus, we did not identify regulators that are specific to some of the remaining inducers. The resources we describe will enable discovery of such regulators in the future. In any case, cell type- and ligand-specific roles of regulators combined with cell type-specific expression are likely a major source of context-dependent AHR signaling. In the future, integrating information on cell type- and ligand-specific roles of regulators with ongoing efforts to identify physiologically relevant ligands and their concentrations in different tissues (83) may enable predictions of which ligands activate AHR, how strongly, and in which cell types and physiological settings. In addition, our approaches can also be used to dissect regulation of AHR activity in the presence of mixtures of inducers, which is particularly relevant for inducers derived from the gut microbiome, and to define mechanisms that give rise to different transcriptional responses in response to different inducers. We expect that such knowledge will be essential to understanding how AHR shapes host-microbe signaling, immune responses, xenobiotic detoxification, and additional physiological processes.

Due to these important roles, AHR is also a sought-after target for therapeutics for inflammatory and metabolic diseases. Although some AHR-targeting compounds have entered the clinic, such as tapinarof for treatment of psoriasis, systemic inhibition or activation of AHR is not a feasible therapeutic strategy because of the resulting effects on additional processes, including xenobiotic metabolism and intestinal barrier integrity.

Our findings may enable manipulation of signaling through targeting of nodes that are only active in specific cell types. AHR activity has also emerged as a major determinant of sensitivity to inhibitors of TIPARP (also known as PARP7) (84, 85), which are under clinical evaluation, alone or in combination with checkpoint inhibitors, for treatment of multiple solid tumors (86-88). This link is in clear agreement with the known role of AHR in transcriptional regulation of TIPARP. Our identification of determinants of AHR activity now may enable prediction of responsiveness of tumors and patient stratification in clinical trials.

More broadly, our findings illustrate how even for a signaling pathway that is simple on paper, layers of regulators can diversify signaling to enable responses that vary by input and cell type. We expect that this pattern will recur in additional chemosensory pathways that, like AHR, integrate multiple inputs and have context-dependent outcomes. Many of these pathways also integrate information from each other, as exemplified by previous findings of crosstalk between AHR signaling and NF-κB and estrogen receptor signaling, among others (10, 12, 13), and our finding that STAT3 signaling stabilizes AHR at the protein level. We expect that systematic genetic approaches will continue to play a critical role in decoding these signaling networks (42-46, 89-92).

## Conclusions

Here, we harness systematic genetic screens to identify hundreds of previously uncharacterized regulators of AHR signaling. These regulators collectively act at every step of AHR signaling, and many act in a manner that is specific to an inducer or cell type. Our findings highlight how AHR signaling is underpinned by a complex regulatory network whose composition varies with context, which likely is critical to the ability of AHR to fulfil a diverse range of physiological roles. In addition, our results lay the groundwork for the identification of factors that contribute to dysregulation of AHR signaling in disease and for the development of therapeutics that manipulate AHR signaling in a targeted manner. Finally, our study further illustrates the power of functional genomics in dissecting complex cellular signaling networks.

## Methods

### General cell culture

HepG2 cells were grown in EMEM (GIBCO) supplemented with 10% (v/v) fetal bovine serum (FBS), 100 units/mL penicillin, and 100 μg/mL streptomycin. HEK293T cells were grown in in DMEM (GIBCO) supplemented with 10% (v/v) fetal bovine serum (FBS), 100 units/mL penicillin, and 100 μg/mL streptomycin. U-87 cells were grown in EMEM (GIBCO) supplemented with 10% (v/v) fetal bovine serum (FBS), 1% (v/v) 1 M HEPES buffer, 100 units/mL penicillin, and 100 μg/mL streptomycin. All parental cell lines were purchased from ATCC.

All cell lines were grown at 37°C in the presence of 5% CO2. All cell lines were periodically tested for Mycoplasma contamination using either the Mycoplasma PCR Detection Kit (Applied Biological Materials Inc) or the Universal Mycoplasma Detection Kit (ATCC).

### Lentivirus production

To generate lentivirus, HEK293T cells were transfected with the transfer plasmid and four packaging plasmids (for expression of VSV-G, Gag/Pol, Rev, and Tat) using TransIT-LT1 Transfection Reagent (Mirus Bio). Viral supernatant was harvested two days post transfection and filtered through 0.44 µM PES filters or frozen at -80°C before transduction.

### Cloning and optimization of AHR reporters

The AHR GFP reporter constructs were cloned into pBA407, a previously described backbone for lentiviral expression (<u>Adamson et al., 2016</u>), by Gibson assembly using the NEBuilder HiFi DNA assembly master mix (NEB). We tested several factors before choosing the optimal reporter including the number of AHR binding sites (4,7,10, and 15 AREs), the spacing between them (18, 23, 51, 70 bp), and the identity of the minimal promoter (cFos, CMV). The final construct contains 15 AREs, spaced by 23 bp stuffer sequences that were randomized to minimize recombination, upstream of the cFos minimal promoter (−53 to +45 of the human c-fos promoter). The resulting vector is pMV0021 (15ARE23bp-GFP, Addgene # 246420).

The luciferase reporter constructs were based on pMV0021. They comprise two expression cassettes that are inverted with respect to the lentiviral backbone to allow transcription of the lentiviral genome to progress normally. The first expression cassette contains 15 AREs, spaced 23 bp apart, upstream of the minimal cFos promoter followed by a firefly luciferase (cloned from Addgene #113088) and the bGH polyA transcription terminator. The second expression cassette contains a UBC constitutive promoter followed by a renilla luciferase (cloned from Addgene #113088), P2A, and eGFP. The renilla luciferase allows for the normalization of firefly luciferase signal by cell number.

In finalizing this construct, we tested two transcription terminators (SV40, BGH), two constitutive promoters (UBC, PGK) and two minimal promoters (p(cFos), p(TATA)). The p(TATA) promoter was cloned using oligos based on the minP sequence in pGL4.23-IL2RA CaRE4 scramble (plus strand) (Addgene #91852). The final vector used for the microbiome chemical screen is pMV0067 (15ARE23bp -p(cFos)-fLuc-UBC-rLuc-P2A-GFP, Addgene # 246421). All subsequent experiments use pHL047 (15ARE23bp - p(TATA)-fLuc-UBC-rLuc-P2A-GFP, Addgene #246422). Both reporters show comparable induction and dynamic range.

### Generation of cell lines

The HepG2 AHR reporter CRISPRi cell lines were constructed in the background of a previously described HepG2 CRISPRi cell line (<u>Replogle et al., 2022</u>) stably expressing Zim3-dCas9-P2A-mCherry under the control of the EF1a promoter.

To make the GFP reporter cell line, the HepG2 CRISPRi cell line was transduced with lentivirus derived from pMV0021 at low multiplicity of infection. This population was treated with 100 µM indole-3-pyruvate for 24 h and a polyclonal population of cells with high GFP was sorted using fluorescence-activated cell sorting (FACS) on a FACSAria II Cell Sorter (BD Biosciences). This polyclonal population was used to generate monoclonal cell lines using limiting dilution. Briefly, the polyclonal cell line was plated in 96-well plates at a ratio of 0.5 cells/well. Cells were grown with fresh media added to each well every ∼3 days. Identified clones were expanded and tested for their level of AHR induction upon treatment with 100 µM indole-3-pyruvate or 100 µM ITE for 24 h. The clonal cell line cMV011 demonstrated uniform and strong induction of GFP fluorescence and was thus used for all GFP experiments.

To generate a U-87 cell line stably expressing Zim3-dCas9, U-87 cells were infected with lentivirus containing UCOE-SFFV-Cas9-P2A-mCherry (pNM1123) (<u>Replogle et al., 2022</u>) at low multiplicity of infection. Polyclonal populations of mCherry+ cells were selected using two rounds of FACS on a FACSMelody™ Cell Sorter (BD Biosciences). A monoclonal GFP reporter cell line (cKD004) was generated using the protocol described above.

To make the luciferase reporter cell lines, the HepG2 CRISPRi cell line was transduced with lentivirus derived from pMV0067 or pHL047 at low multiplicities of infection. Polyclonal populations of GFP+ cells were selected using FACS on a FACSAria II Cell Sorter (BD Biosciences). The resulting polyclonal cell lines (cMV007, cMV023) were used for luciferase experiments.

### Genome-wide CRISPRi screen

FACS-based, genome-scale CRISPRi screens were conducted using protocols similar to those previously described (<u>Gilbert et al., 2014; Horlbeck et al., 2016; Sidrauski et al., 2015</u>, <u>Adamson et al., 2016</u>; <u>Hickey at al., 2020</u>). The human CRISPRi-v2 top 5 (Addgene, Cat#83969) sgRNA library was transduced into cMV011 cells at an MOI < 1 in duplicate (BFP+ cell percentages were ∼39% and 28%, respectively). Replicates were carried separately throughout the experiment. The cells were grown for 3 days without selection, followed by 5-7 days of selection with 1-2 µg/mL puromycin. The cells were removed from selection for 1 day and seeded for treatment. Cells from both replicates were treated with 100 µM indoxyl sulfate for 24 h. A separate batch of cells from one of the replicates was treated with 100 µM indole-3-pyruvate for 24 h. After 24 h of treatment, cells with the highest and lowest GFP fluorescence (25-30%) were isolated by FACS on a FACSMelody™ Cell Sorter (BD Biosciences). Approximately 25-30 million cells were collected per bin. Cells were washed with PBS and cell pellets were frozen at −80°C after collection.

Genomic DNA was isolated from frozen cells using the Macherey Nagel NucleoSpin Blood XL or L kits depending on pellet size. sgRNA-encoding regions were prepared for sequencing using a protocol that omitted previously used gel extraction steps (<u>Replogle et al., 2022</u>). Sequencing was performed on a NextSeq 1000 (Illumina) with custom sequencing primers.

Sequenced protospacer sequences were aligned and data were processed as described (<u>Gilbert et al., 2014; Horlbeck et al., 2016</u>) with custom Python scripts (ScreenProcessing, available at https://github.com/mhorlbeck/ScreenProcessing). Reporter phenotypes for sgRNAs were calculated as the log_2_ ratio of sgRNA counts within the high GFP cells over the low GFP cells, normalized to the median log_2_ ratio of negative control sgRNAs. Phenotypes for each transcription start site (“AHR activation phenotypes”) were then calculated as the average reporter phenotype of the 3 sgRNAs with the strongest phenotype by absolute value (most active sgRNAs). Mann-Whitney test p values were calculated by comparing all sgRNAs targeting a given TSS to the full set of negative control sgRNAs. Screen hits were defined as those genes with a discriminant score > 7, defined as the absolute value of the reporter phenotype over the standard deviation of negative control reporter phenotypes multiplied by the –log_10_ of the Mann-Whitney p value.

### Sublibrary CRISPRi screens

To select genes for the sublibrary screens, we first compiled the union of genes that were hits with either indoxyl sulfate or indole-3-pyruvate in the genome-wide screens (771 genes). We excluded a number of genes involved in central cellular processes, such as ribosomal and proteasomal genes. To this set, we supplemented 68 genes of interest that had absolute phenotype scores > 0.5 in at least one replicate but did not score as hits. Finally, we curated a set of 66 genes that we hypothesized to play a role in AHR activation by other inducers, including transporters, Trp metabolism enzymes, and transcriptional coregulators. The final library targets 497 genes (543 TSSs).

Oligos encoding sgRNA sequences for the 2715 targeting (5 sgRNAs per TSS) and 150 non-targeting sgRNAs were selected from the human CRISPRi-v2 top 5 library and ordered as a pool from Twist Biosciences. The sgRNA sublibrary was cloned into pCRISPRia-v2 (Addgene, Cat# 84832) using protocols similar those previously described (Horlbeck et al., 2016; Heo et al., 2024) and subsequently packaged in lentivirus and titrated.

The FACS-based sublibrary screens were performed similarly to the genome-wide screens. The sublibrary was transduced into cMV011 cells at an MOI < 1 in duplicate (BFP+ cell percentages were ∼26% and 31%, for the two transduction replicates). Screen replicates were carried separately throughout the remainder of the experiment. The cells were grown for 2 days without selection, followed by 8 days of selection with 1 µg/mL puromycin. The cells were allowed to recover from selection for 2 days before seeding for treatment. They were treated for 24 h with one of 14 compounds at the concentrations indicated: 3-methylcholanthrene (10 µM, MilliporeSigma), benzo[a]pyrene (50 µM, MilliporeSigma), ITE (1 µM, Fisher Scientific), indirubin (10 µM, MilliporeSigma), indole-3-acetic acid (100 µM, Sigma-Aldrich), indole-3-carbinol (100 µM, Sigma-Aldrich), indole-3-carboxaldehyde (100 µM, Sigma-Aldrich), indole-3-lactate (200 µM, Sigma-Aldrich), indole-3-pyruvate (100 µM, Sigma-Aldrich), indoxyl sulfate (100 µM, Sigma-Aldrich), kynurenine (100 µM, Sigma-Aldrich), urolithin B (100 µM, Sigma-Aldrich), urolithin M5 (100 µM, Toronto Research Chemicals Inc.), delphinidin (100 µM, Cayman Chemical Company). After 24 h of treatment, cells with the highest and lowest GFP fluorescence (25-30%) were isolated by FACS on a FACSAria II Cell Sorter (BD Biosciences). Approximately 1 million cells were collected per bin. Cells were washed with PBS and cell pellets were frozen at −80°C after collection. For the growth screen, cells were collected once puromycin selection was complete (t0) and after 10 additional doublings (t10).

Genomic DNA was isolated from frozen cells using the Macherey Nagel NucleoSpin Blood Mini kit and the sgRNA-encoding regions were amplified and prepared for sequencing as described above. For the reporter screens, sequencing data was processed, and hits called in the same way as for the genome-wide screen described above, with one distinction: we modified the analysis to require a single TSS to be used to calculate phenotypes across all inducers. We chose the TSS that was a hit with the greatest number of inducers.

For the growth screen, sgRNA phenotypes were calculated as the log_2_ ratio of sgRNA counts at the t10 timepoint over the t0 timepoint, normalized to the median log_2_ ratio of negative control sgRNAs and divided by the number of cell doublings (<u>Kampmann et al., 2013</u>; <u>Gilbert et al., 2014</u>). Phenotypes were calculated as done for the genome-scale screen.

### Chemical screening

Microbiome library screen: Two replicates of cMV007 AHR luciferase reporter cells were seeded in 96-well plates overnight before treatment with the microbiome compound library for 4 h. Each plate had a set of vehicle controls as well as ITE and FICZ as positive controls. The Promega Dual-Luciferase® Reporter Assay System (cat # E1910) was used to conduct luciferase assays for AHR activity using the manufacturer’s protocol. Luminescence was measured on a Tecan Infinite M Plex plate reader (Supplementary Table 3).

Urolithin library screen: Two replicates of cMV023 AHR luciferase reporter cells were seeded in 96-well plates overnight before 4 h treatment with 1, 10, or 100 µM of each compound in the urolithin library. The plate had a set of vehicle controls as well as ITE as a positive control. Luciferase assay was performed as before, and luminescence was measured on a Perkin Elmer EnSight plate reader (Supplementary Table 4).

Fold change in AHR induction for each compound was calculated by dividing the ratio of firefly to renilla RLUs by corresponding ratios for the vehicle controls on the same 96-well plate. Compound toxicity was assessed using % mean renilla RLU of all conditions as a proxy for relative cell number between treatments. Samples with % mean renilla RLU ≤ 50% were excluded.

### Individual sgRNA retests

To independently validate the genome-wide CRISPRi screen, a subset of hits was selected. The best performing guide for each gene based on the screen phenotypes was cloned into pCRISPRia-v2 (Addgene, Cat#84832) and packaged into lentivirus as described previously (Horlbeck et al., 2016, <u>Jost et al., 2017</u>). cMV011 cells were seeded and transduced with the sgRNA of interest at MOI <1. Transduced cells were allowed to grow for ∼10 days and treated with compound of interest for 24 h and GFP induction was measured via flow cytometry.

Growth phenotypes for each sgRNA were measured as the log_2_ ratio of the ratio of sgRNA-positive cells to sgRNA-negative cells at 14 days post-transduction compared to 2 days post-transduction, normalized by the equivalent ratio for the non-targeting control sgRNA and divided by number of days. sgRNAs with < 100 sgRNA-expressing cells recorded were excluded from further analysis. For the remaining sgRNAs, a phenotype score was calculated as log_2_[(median GFP of sgRNA+ cells)/(median GFP of sgRNA-cells)] without normalizing for autofluorescence. To calculate % knockdown, cells were selected with 1-2 µg/ml puromycin for 7-10 days. RNA was extracted and level of knockdown was measured via RT-qPCR using primers specific to the gene of interest as outlined below.

To assess effects of general regulators in U-87 cells, a similar protocol as described above was followed using cKD004 U-87 AHR-GFP reporter CRISPRi cells. To measure transcript levels of AHR, ARNT, or TIPARP or AHR protein levels when general regulators are perturbed, HepG2 CRISPRi cells were transduced with sgRNAs targeting general regulators as before and selected with 1-2 µg/ml puromycin before seeding for RNA and protein extraction.

All sgRNA sequences used for individual retests are listed in Supplementary Table 12.

### RNA Extraction and RT-qPCR

Cells were washed with PBS and stored at −80°C until RNA extraction. Total RNA was extracted using the NEB Monarch® Total RNA Miniprep Kit (Catalog # T2010) as per the manufacturer’s instructions. RT-qPCR analysis was conducted using the Luna® Universal One-Step RT-qPCR Kit (NEB, Catalog # E3005) on an Applied Biosystems QuantStudio 6 Pro real-time PCR 384-well system. Fold change values were calculated using the 2^−ΔΔCt^ method, with β-actin as the housekeeping gene.

The primer sequences are listed in Supplementary Table 13.

### Western blotting

Cells were lysed using 1X RIPA buffer (Sigma #20-188) supplemented with a protease inhibitor cocktail (Sigma #P8340). Protein concentration was determined using the Pierce™ BCA Protein Assay Kits (Thermo Scientific #23225) and normalized. Protein samples were denatured with SDS and ∼10 µg of protein was separated on a 10% TGX gel (BioRad #4561036). Proteins were transferred to a PVDF membrane (Thermo Scientific #88518) using wet transfer electrophoresis. Membranes were blocked in 5% non-fat milk in TBST and probed with appropriate primary (AHR: 83200, ACTB: 4970, GAPDH: 5174, α-tubulin: 2125, Cell Signaling Technology; AIP: ab228684, Abcam) and secondary (anti-rabbit horseradish peroxidase-conjugated secondary antibody, 7074, Cell Signaling Technology) antibodies. The membrane was incubated in Clarity Max Western ECL Blotting Substrate (BioRad, #1705062) and proteins bound by both primary and secondary antibodies were visualized by chemiluminescence on a ChemiDoc gel imaging system (BioRad). Proteins were quantified by normalizing the intensity of the protein band of interest to the intensity of a housekeeping gene (GAPDH, ACTB/β-actin, or α-tubulin) band in the same lane using Fiji software.

## Acknowledgements

We thank Hyunho Lee for sharing pHL047, the p(TATA) luciferase vector, Kelsey Hickey for sharing neddylation and proteasome inhibitors, Deepsing Syangtan for preparing the urolithin chemical library, and Carol Gross, Jonathan Weissman, Max Horlbeck, and all members of the Jost lab for helpful discussions. pBA407 was a gift from Jonathan Weissman (Addgene plasmid # 85970; http://n2t.net/addgene:85970; RRID:Addgene_85970). pBA409 was a gift from Jonathan Weissman (Addgene plasmid # 85971; http://n2t.net/addgene:85971; RRID:Addgene_85971). PAK4-FL-3’UTR was a gift from James Donahue (Addgene plasmid # 113088; http://n2t.net/addgene:113088; RRID:Addgene_113088). pGL4.23-IL2RA CaRE4 scramble (plus strand) was a gift from Jacob Corn (Addgene plasmid # 91852; http://n2t.net/addgene:91852; RRID:Addgene_91852). We thank the Harvard Immunology Flow Cytometry Core Facility for access to FACS machines and flow cytometers as well as the BPF NGS Genomics Core Facility for access to Tapestation and Bioanalyzer machines. This work was supported in part by the National Institutes of Health (grants R00GM130964 and DP2GM154152 to MJ, K08AI130392-01, DP2GM136652, and R01AI1269151 to SRN), a Charles H. Hood Foundation Child Health Research Award (MJ), the G. Harold and Leila Y. Mathers Foundation (MJ), the William F. Milton Fund at Harvard University (MJ), a Springer-Nature Global Grant for Gut Health (MJ), a Career Award for Medical Scientists from the Burroughs Wellcome Fund (SRN), a Pew Biomedical Scholarship (SRN), and the Food Allergy Science Initiative (SRN). EPB is a Howard Hughes Medical Institute Investigator. KD was supported by the EPSRC Centre for Doctoral Training in BioDesign Engineering (EP/S022856/1).

## Author Contributions

MV performed all experiments, analyzed data, and wrote the manuscript. KD constructed the U-87 CRISPRi cell line and the U-87 AHR reporter cell line. YD, XW, MAF, and MJ curated and compiled the microbiome chemical library. MB and EPB designed and compiled the library of urolithins and related chemicals. SRN contributed to conceptualization. MJ conceived and supervised experiments and wrote the manuscript. All authors provided feedback on the manuscript.

## Supplemental figures and figure legends

**Figure S1.**
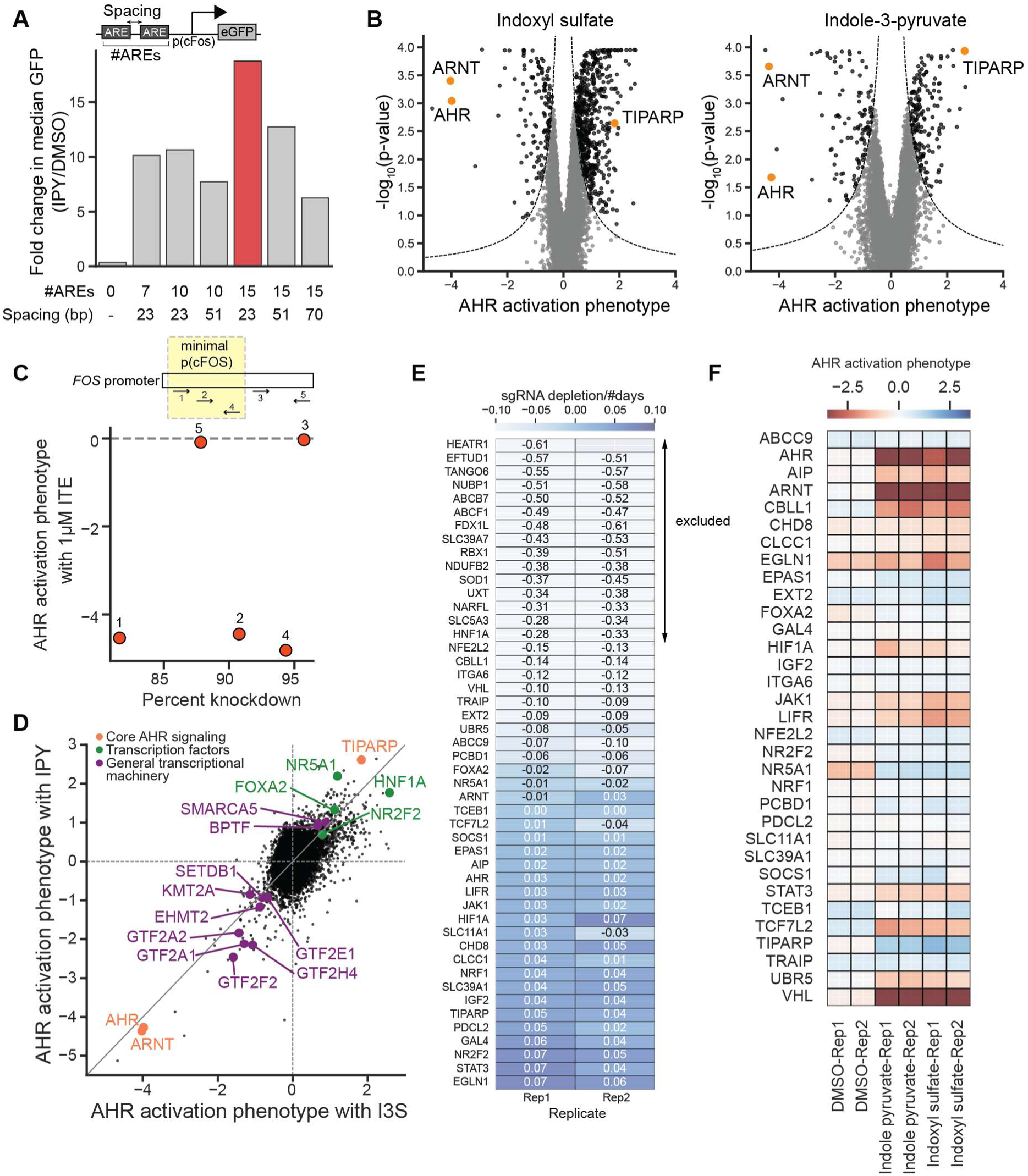
Genome-scale CRISPRi screens reveal modulators of AHR signaling. (A) Schematic of GFP reporter designs showing the number of AHR response elements (AREs) and spacing between consecutive AREs, and fold change in GFP signal upon induction with 100 µM IPY for 24 h. The final reporter is highlighted in red. (B) Volcano plots for the genome-scale screens for AHR regulators with I3S and IPY. Three positive control genes, AHR, ARNT and TIPARP, are highlighted. Dashed line shows the threshold used for calling hits; gray points are not hits, black points are hits. (C) Top: schematic representation of FOS promoter with binding sites for the 5 sgRNAs targeting FOS indicating the p(cFOS) promoter used in the reporter. Bottom: effect of each of the 5 sgRNAs on AHR activation by 1 µM ITE plotted against the percent knockdown of the FOS gene measured by RT-qPCR. (D) AHR activation phenotypes for all genes in the IPY and I3S CRISPRi screens. Categories of genes highlighted include core AHR signaling genes (orange), other transcription factors (green), and genes that are components of the general transcriptional machinery (purple). (E) Depletion of sgRNA-expressing cells over the 14-day period of the validation experiment as a proxy for gene essentiality. 15 genes excluded from analysis are highlighted. (F) AHR activation phenotypes calculated using the log_2_ ratio of median GFP fluorescence in sgRNA-expressing cells relative to median GFP fluorescence for non-sgRNA-expressing cells in the same well after treatment with 100 µM I3S, 100 µM IPY, or vehicle control for 24 h.

**Figure S2.**
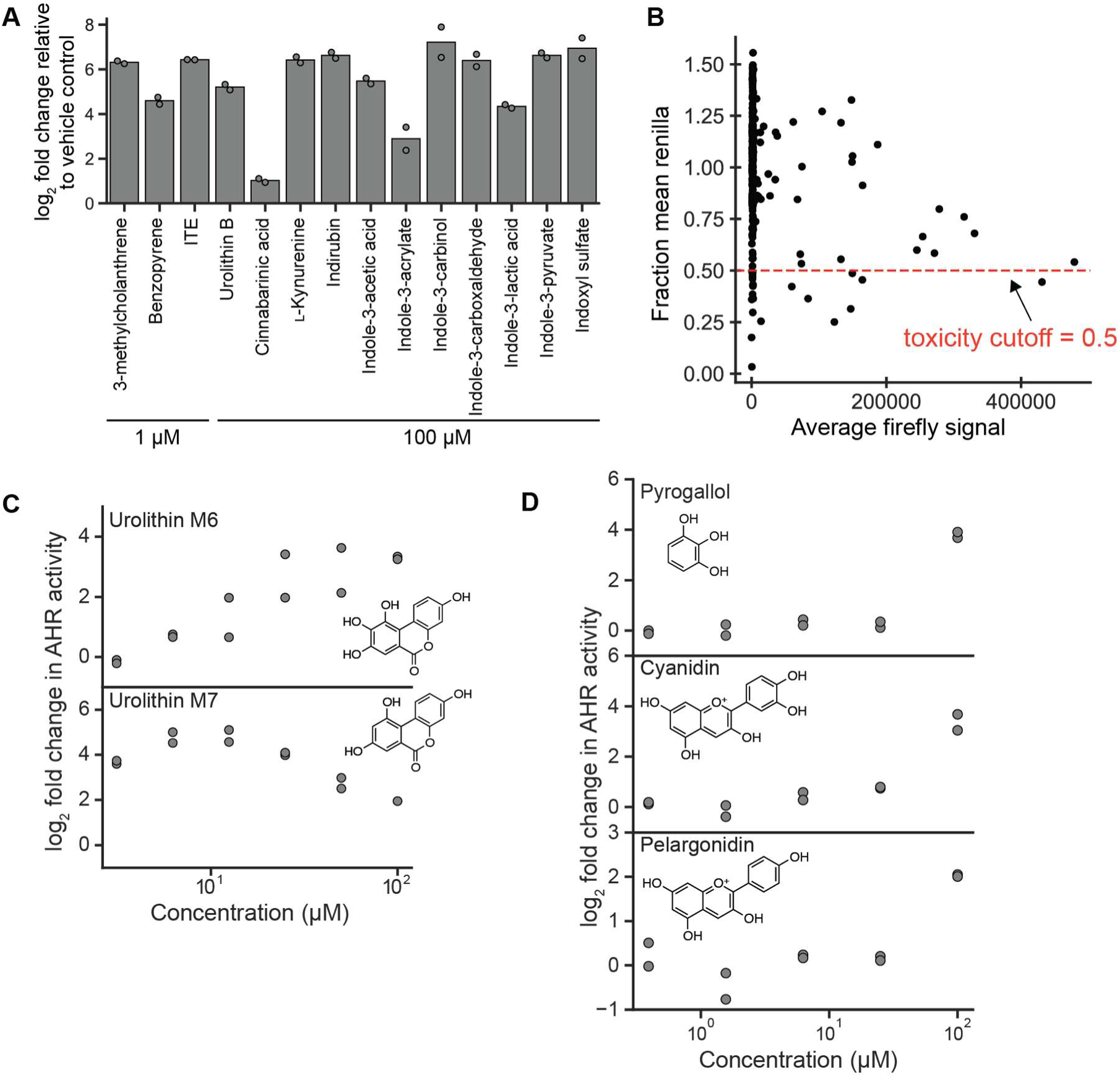
Chemical screens identify novel AHR inducers. (A) log_2_ fold change in luciferase signal after treatment with the indicated chemicals for 4 h relative to vehicle control. (B) Renilla relative luminescence units (RLUs, average of duplicate treatments) normalized to mean Renilla RLUs across the entire library plotted against firefly RLUs (average of duplicate treatments) for all chemicals in the 250-chemical library. All chemicals with fraction mean Renilla RLUs ≤ 0.5 are excluded from analysis. (C) Dose-response curves of the two additional urolithins that activate AHR, after treatment of the HepG2 luciferase reporter cells for 4 h. (D) Dose-response curves of 3 additional chemicals from the 250-chemical screen that induce AHR activation, after treatment of the HepG2 luciferase reporter cells for 4 h. Data represent treatment replicates (*n*=2).

**Figure S3.**
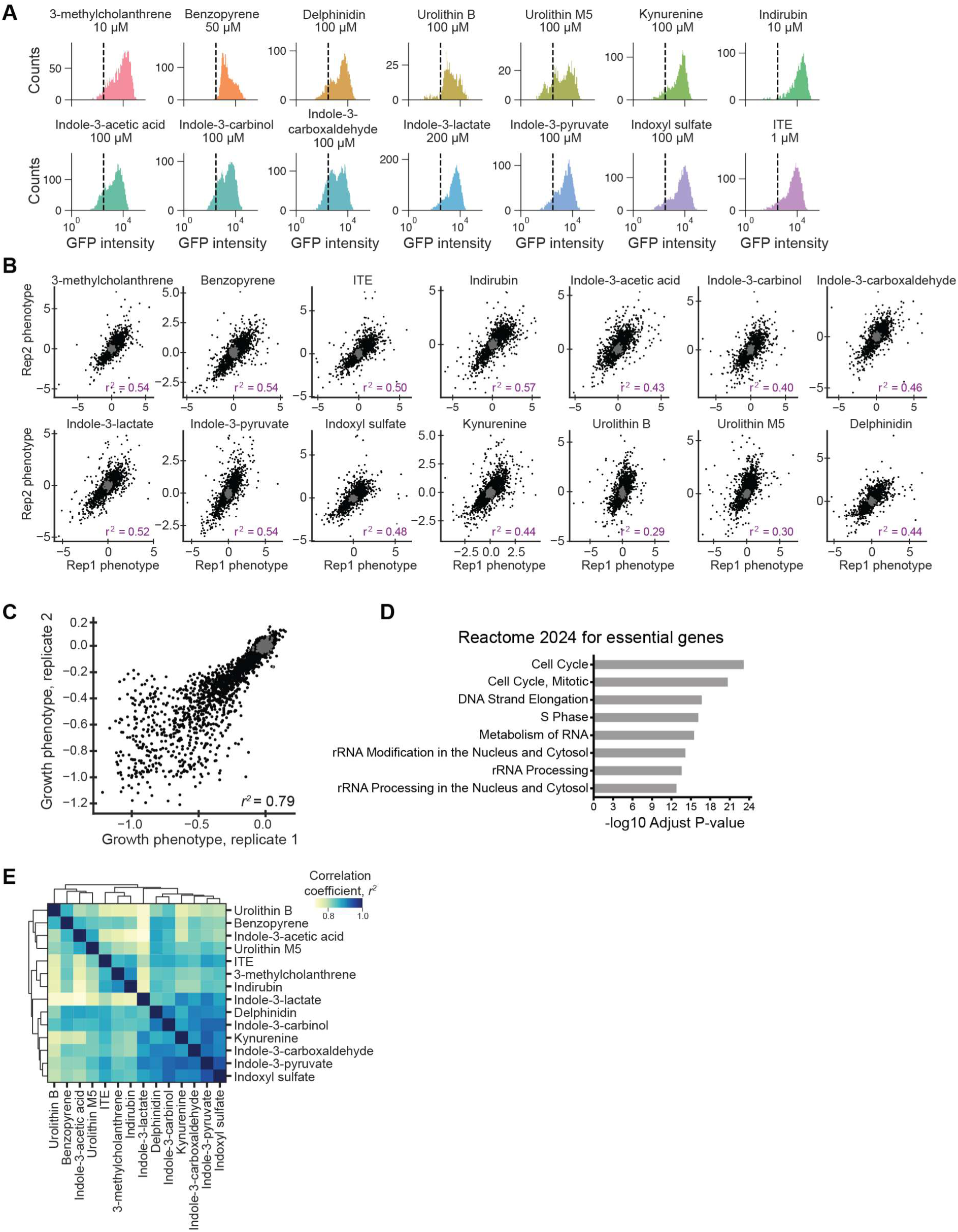
Sublibrary CRISPRi screens reveal AHR regulatory networks across inducers. (A) GFP fluorescence distributions of the monoclonal CRISPRi GFP reporter cell line (cMV011) treated with indicated concentrations of the 14 chemicals used in the sublibrary screens. Dashed line indicates the median GFP fluorescence for vehicle-treated cells. (B) Replicate correlation for the FACS-based screens with all inducers for all targeting sgRNAs (black) and non-targeting control sgRNAs (gray). (C) Growth phenotypes from two biological replicates for all targeting sgRNAs (black) and non-targeting control sgRNAs (gray). (D) Gene set enrichment for all genes identified to be essential from the growth screen, using the Reactome 2024 dataset. (E) Pairwise correlations (squared Pearson correlation coefficient, *r*^2^) of gene-level AHR activation phenotypes for all inducers. Phenotype sets were clustered using Euclidean distance as indicated by the dendrograms.

**Figure S4.**
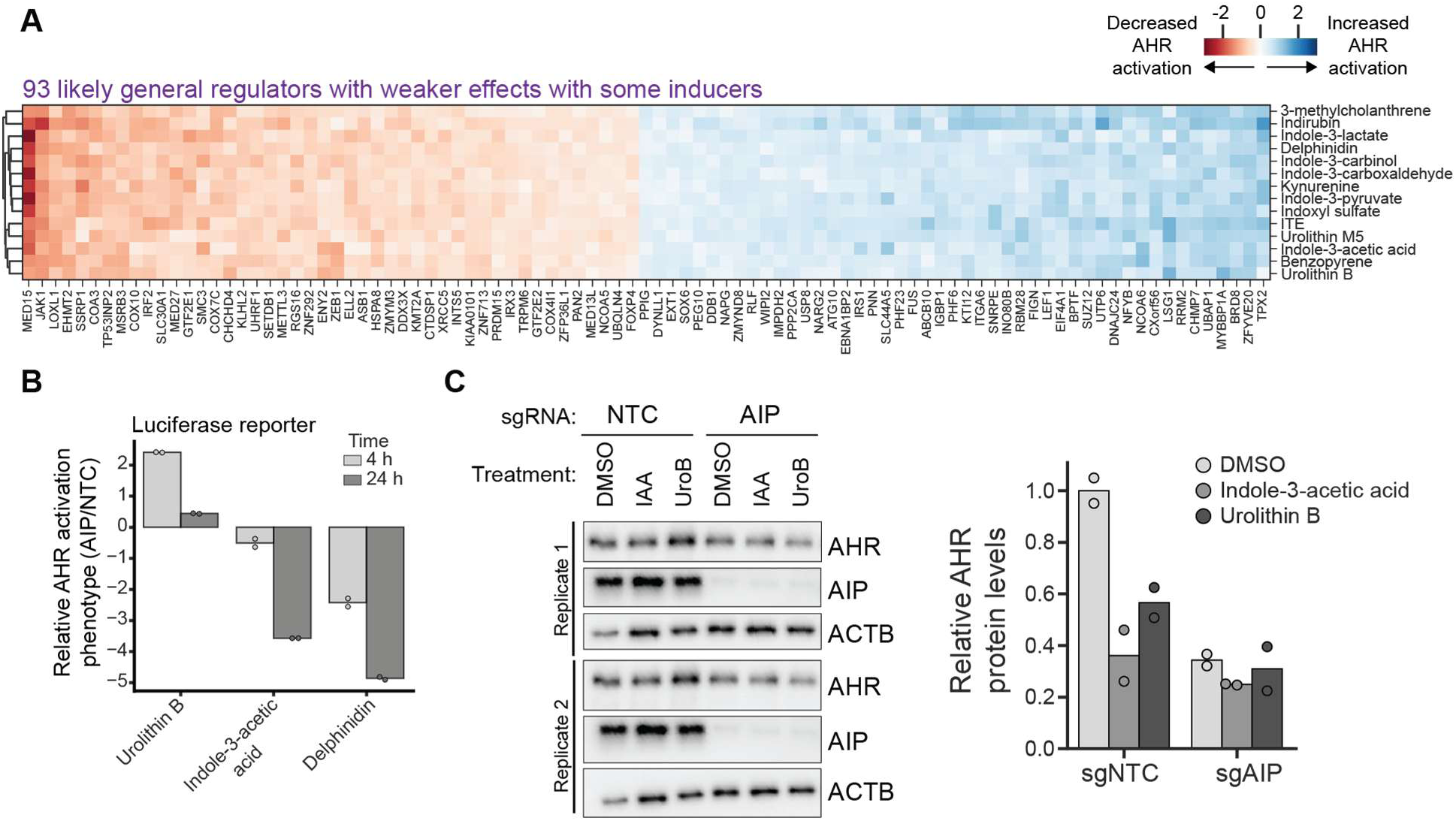
Identification of general and inducer-specific regulators of AHR signaling. (A) AHR activation phenotypes for the 93 genes that are likely general regulators but have weaker effects with some inducers. (B) AHR activation phenotypes for AIP knockdown from HepG2 luciferase reporter cells. AHR luciferase reporter cells were transduced with either a non-targeting control sgRNA or a sgRNA targeting AIP, selected for sgRNA-expressing cells, and treated with 100 µM of the indicated chemicals for either 4 or 24 h. AHR activation phenotypes of AIP knockdown cells were normalized to phenotypes of cells with a non-targeting control sgRNA. (C) Left: Western blots for AHR and AIP for HepG2 CRISPRi cells transduced with either a non-targeting control or an sgRNA targeting AIP before 4 h treatment with 100 µM indole-3-acetic acid (IAA), 100 µM urolithin B (UroB), or vehicle control. Right: Quantification of AHR protein levels relative to the beta-actin loading control normalized to the negative control (sgNTC, vehicle control). The decrease in AHR protein levels upon treatment with either inducer in non-targeting control cells is expected due to known degradation of ligand-bound AHR after nuclear translocation and DNA binding.

**Figure S5.**
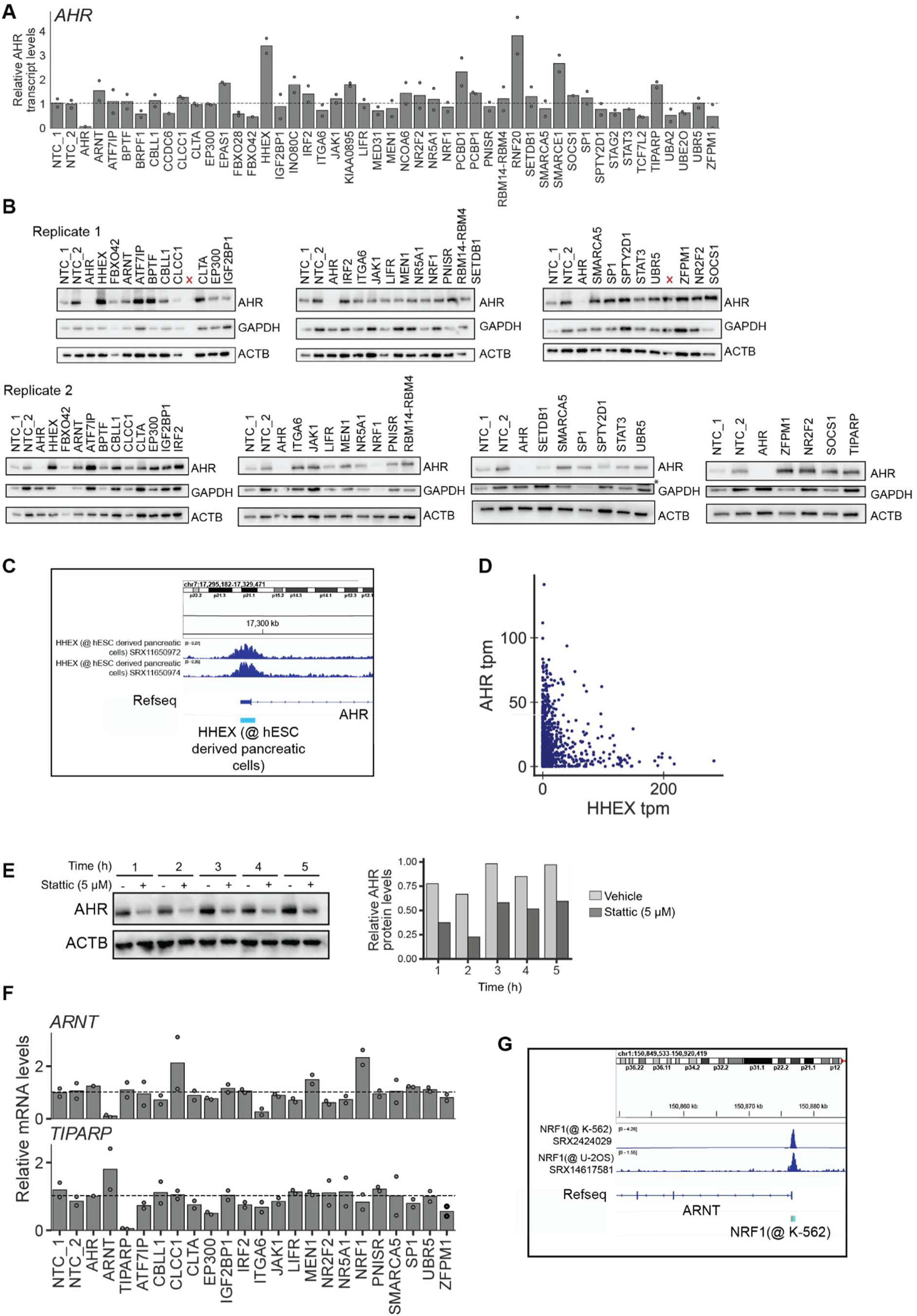
Functional classification of general regulators of AHR signaling. (A) AHR mRNA levels for all knockdown cells relative to the average level for cells with the two non-targeting control sgRNAs, shown by the dashed line, as measured using RT-qPCR. (B) Western blots for AHR and two loading controls of HepG2 CRISPRi cells transduced with sgRNAs targeting the general regulators not classified as regulators of AHR mRNA levels. The red crosses indicate lanes that were empty or re-loaded due to broken wells. The asterisk (*) denotes a lane with a non-specific band that appears due to incomplete stripping of ACTB before probing with the GAPDH antibody. (C) ChIP-seq tracks for HHEX in ESC-derived pancreatic cells (71) upstream of the AHR gene. (D) AHR and HHEX transcript per million (tpm) for all cell lines in the Human Protein Atlas. (E) Left: Western blots for AHR of HepG2 cells treated with 5 µM stattic, a STAT3 inhibitor, for the indicated times. Right: Quantification of AHR relative to beta-actin loading control. (F) ARNT (top) and TIPARP (bottom) mRNA levels for HepG2 cells transduced with sgRNAs against the indicated genes relative to the average levels for cells with the two non-targeting sgRNAs, indicated by the dashed line, as measured using RT-qPCR. (G) ChIP-seq tracks of NRF1 upstream of the ARNT gene from the ENCODE database in K-562 chronic myeloid leukemia cells and U-2OS osteosarcoma cells.

**Figure S6.**
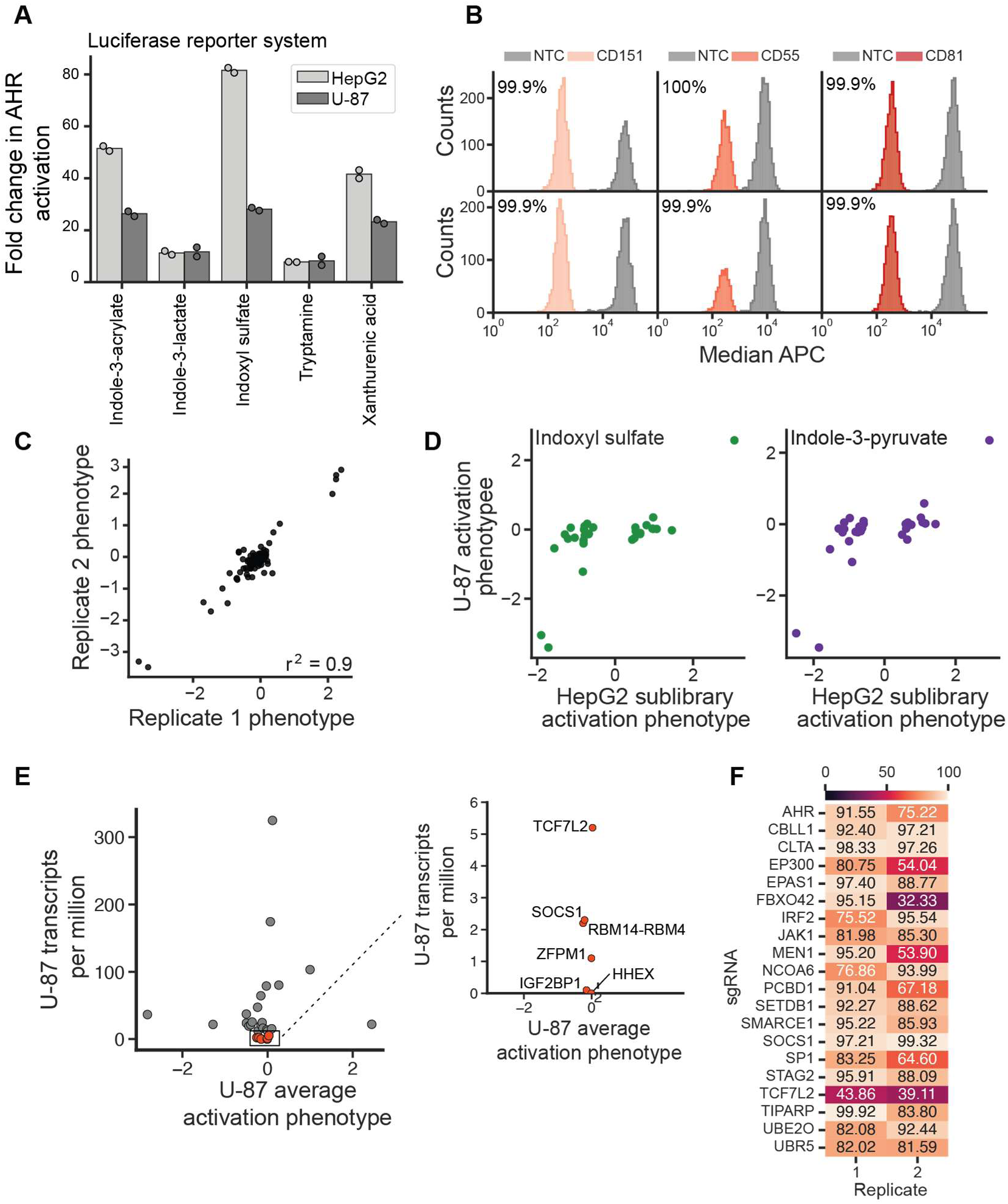
Many general regulators of AHR signaling act in a cell type-specific manner. (A) Fold change in AHR luciferase reporter signal upon treatment with 100 µM of the indicated molecules for 4 h in HepG2 and U-87 cells. (B) Distribution of anti-CD151, anti-CD55, and anti-CD81 signal intensities (APC) in cells with a targeting sgRNA or a non-targeting control sgRNA, for two independently transduced replicates. Knockdown was calculated as the ratio of median APC signal in cells transduced with targeting sgRNAs relative to cells transduced with a non-targeting control sgRNA. (C) Replicate correlation of AHR activation phenotypes for U-87 cells transduced with sgRNAs targeting 27 general regulators, 3 positive control sgRNAs and 2 non-targeting control sgRNAs. (D) Comparison of AHR activation phenotypes for U-87 and HepG2 cells for the selected sgRNAs for I3S and IPY. (E) Transcripts per million of targeted genes in U-87 cells from the Human Protein Atlas plotted against the U-87 AHR activation phenotypes averaged for I3S and IPY. Red points and inset indicate genes with < 6 transcript per million. (F) % knockdown for a subset of genes in the U-87 cells for two independently transduced replicates, measured using RT-qPCR.

**Figure S7.**
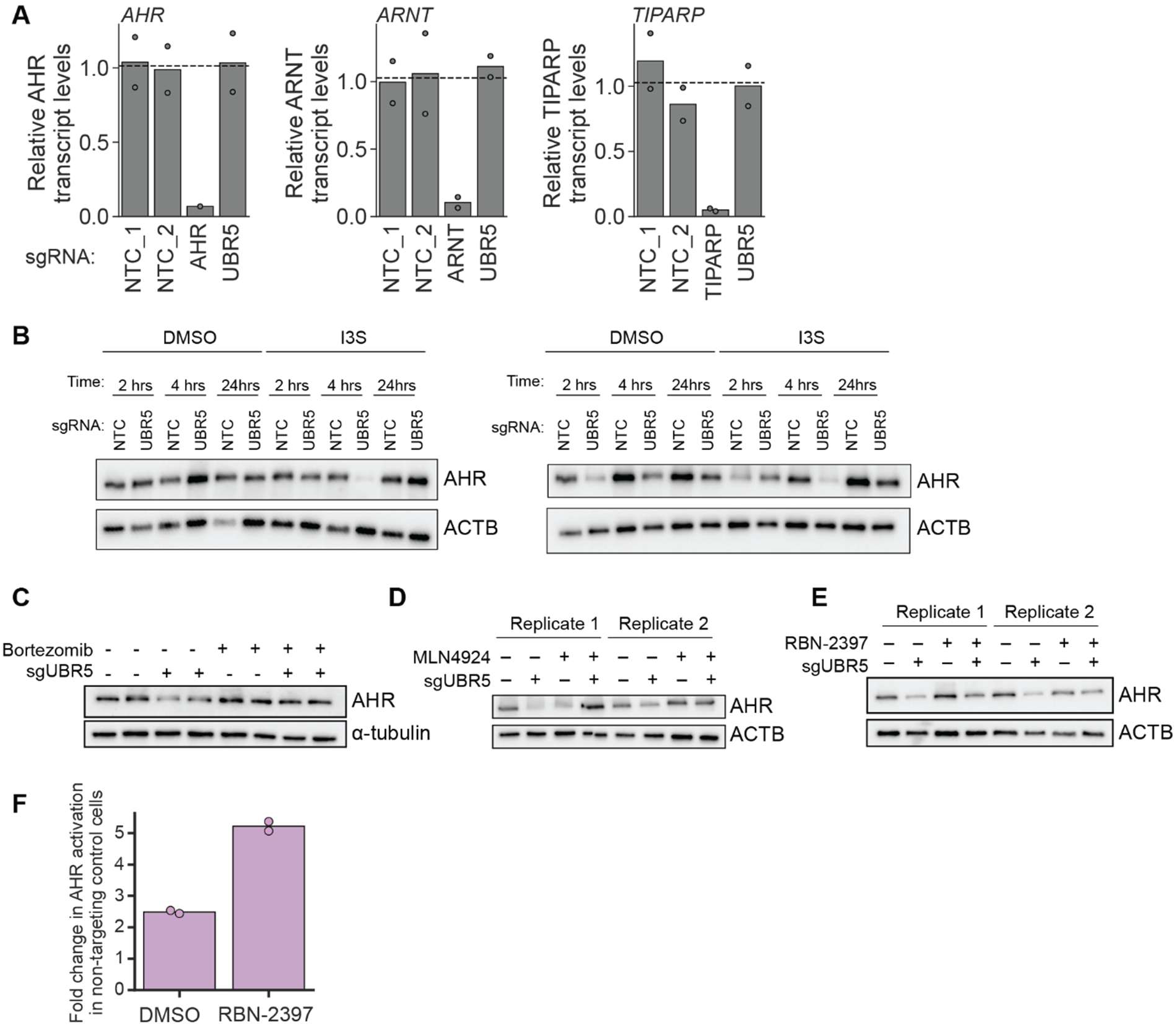
The E3 ubiquitin ligase UBR5 is a regulator of AHR turnover. (A) Relative mRNA levels for AHR, ARNT, and TIPARP in HepG2 CRISPRi cells transduced with two non-targeting control sgRNAs, 1 positive control sgRNA, or an sgRNA targeting UBR5. (B) Western blots for AHR of HepG2 cells transduced with an sgRNA targeting UBR5 or a non-targeting control sgRNA, treated with 100 µM I3S for the indicated times. Blots indicate duplicate transductions. (C) Western blots of UBR5 knockdown or non-targeting control cells treated with 100 µM IPY for 4 h after a 2 h pretreatment with 500 nM bortezomib or vehicle control. Blots indicate duplicate treatments. (D) Western blots of UBR5 knockdown or non-targeting control cells treated with 100 µM I3S for 4 h after a 2 h pretreatment with 1 µM MLN4924 or vehicle control. Blots indicate duplicate transductions. (E) Western blots of UBR5 knockdown or non-targeting control cells treated with 100 µM I3S for 4 h after a 4 h pretreatment with 1 µM RBN-2397 or vehicle control. Blots indicate duplicate transductions. (F) Fold change in AHR activation by 100 µM IPY relative to vehicle control in HepG2 cells transduced in duplicate with a non-targeting control sgRNA.

## Supplementary Table List

Table S1. sgRNA counts and phenotypes for genome-scale CRISPRi screen for AHR activation

Table S2. Gene-level phenotypes for genome-scale CRISPRi screen for AHR activation

Table S3. Raw data and calculated fold changes for AHR activation by microbiome chemical library

Table S4. Raw data and calculated fold changes for AHR activation by urolithin and related chemicals

Table S5. sgRNA counts for sublibrary CRISPRi screen for AHR activation

Table S6. sgRNA phenotypes for sublibrary CRISPRi screen for AHR activation

Table S7. Gene-level phenotypes for sublibrary CRISPRi screen for AHR activation

Table S8. sgRNA counts for sublibrary CRISPRi growth screens

Table S9. sgRNA phenotypes for sublibrary CRISPRi growth screens

Table S10. Gene-level phenotypes for sublibrary CRISPRi growth screens

Table S11. Classification of genes based on their AHR activation phenotypes across inducers

Table S12. sgRNA sequences used for individual retests Table S13. RT-qPCR primer sequences

